# The structure of the Orm2-containing serine palmitoyltransferase complex reveals distinct inhibitory potentials of yeast Orm proteins

**DOI:** 10.1101/2024.01.30.577963

**Authors:** Carolin Körner, Jan-Hannes Schäfer, Bianca M. Esch, Kristian Parey, Stefan Walter, David Teis, Dovile Januliene, Oliver Schmidt, Arne Moeller, Florian Fröhlich

**Author notes:** These authors contributed equally to this work (Carolin Körner, Jan-Hannes Schäfer). Correspondence (F.F.), (A.M.), (O.S.).

## Abstract

The levels of sphingolipids are crucial determinants of neurodegenerative disorders. Therefore, sphingolipid levels have to be tightly regulated. The Orm protein family and ceramides act as inhibitors in the rate-limiting step of sphingolipid biosynthesis – the condensation of L-serine and palmitoyl-CoA. The two yeast isoforms Orm1 and Orm2 form a complex with the serine palmitoyltransferase (SPT). While the sequences of the Orm proteins are highly similar, however, Orm1 and Orm2 are differentially regulated in yeast cells. The mechanistic details of the differential regulation remain elusive. To elucidate the regulatory mechanism, we determined the cryo-electron microscopy structure of the SPT complex containing Orm2. Through *in vitro* activity assays and a newly developed plasmid coverage assay combined with targeted lipidomics, we demonstrate that the SPT complex with Orm2 exhibits lower activity in comparison to the complex containing Orm1. This collectively suggests a higher inhibitory potential of Orm2, despite the remarkably similar structures of the Orm1- and Orm2 containing complexes. The high conservation of the SPT from yeast to man implies that the three human ORMDL isoforms could also have different regulatory capacities, which might be essential to understanding their role in sphingolipid-mediated neurodegenerative disorders.

**Highlights:**

- Cryo-EM structures of serine palmitoyltransferase with Orm2 from *S. cerevisiae*
- Ceramide-mediated inhibition on SPT activity is conserved amongst SPTs
- The two yeast Orm isoforms have different regulatory capacity

## Introduction

Sphingolipids are highly enriched in the central nervous system (CNS) and play a pivotal role in neuronal function. Alterations in sphingolipid levels can lead to loss of synaptic plasticity, cell death, and neurodegeneration and have been linked to several neurological diseases^1^. Mutations in the lysosomal enzymes glucocerebrosidase (GBA1) and acid sphingomyelinase (SMPD1) that are required for sphingolipid degradation result in lysosomal storage diseases such as Gaucher disease (GBA) and Niemann-Pick disease type A/B^2–4^. In contrast, hereditary sensory and autonomic neuropathy type 1 (HSAN1) and childhood-onset amyotrophic lateral sclerosis (ALS) are caused by mutations in sphingolipid biosynthesis genes encoding subunits of the serine palmitoyltransferase (SPT)^5–8^.

At the endoplasmic reticulum (ER), SPT catalyzes the first and rate-limiting step in sphingolipid biosynthesis, which is the pyridoxal 5-phosphate (PLP)-dependent condensation of L-serine and palmitoyl-CoA to 3-ketosphinganine (3-KS)^9^. 3-KS is then converted into long chain bases (LCBs), which are further processed to ceramides and complex sphingolipids^10^. The SPT enzyme is remarkably conserved across species and consists of two large catalytic subunits (Lcb1 and Lcb2 in yeast; SPTLC1 and SPTLC2/3 in humans) and a small regulatory subunit (Tsc3 in yeast; SPTssA/B in humans)^11–17^. Orm proteins (Orm1 and Orm2 in yeast; ORMDL1-3 in human) act as negative regulators, inhibiting SPT activity in response to elevated sphingolipid levels^18,19^. In yeast, the phosphatidylinositol-4-phosphate (PI4P) phosphatase Sac1 further associates with the SPT (known collectively as the SPOTS complex) and has been shown to modulate SPT activity^19–21^.

We have recently solved the structure of the yeast SPT as monomers and dimers in complex with its negative regulator Orm1. Our data revealed that Sac1 selectively interacts with Lcb2 of the monomeric SPOTS complex^21^. While the regulatory subunit Tsc3 interacts with Lcb2 to promote SPT activity, Sac1 inhibits SPT activity ^16,19,21–23^. Previous cryo-EM studies of the human SPT-ORMDL3 and *A. thaliana* SPT-ORM1 complexes revealed dimers with a characteristic crossover of the individual subcomplexes^24–26^. Importantly, the predominant form of the endogenous SPOTS complex in yeast appears to be the monomeric form, as indicated by co-immunoprecipitations. The structure of the dimeric yeast SPOT-Orm1 complex did not show an equivalent helix-crossover of Lcb1 as observed in other species^21^. Furthermore, our findings provided mechanistic insights into the homeostatic regulation of SPT activity by binding its downstream metabolite ceramide. Ceramide is complexed at the interface between Orm1 and Lcb2, locking the SPT complex in an inhibited state. The SPT-Orm1 complex thus probably acts as an intra-membrane ceramide-sensor^21^. This ceramide-mediated inhibitory effect on SPT appears to be evolutionarily conserved, although, in higher eukaryotes, ceramide was rather suggested to exert its inhibitory function by stabilizing the N-terminus of ORM/ORMDL proteins that affects SPT activity^26–30^.

While ceramides and ORM proteins inhibit SPT activity, the upregulation of SPT activity (and the ensuing increase in sphingolipid biosynthesis) in yeast is mediated through phosphorylation of amino-terminal residues of the Orm proteins by TORC2/Ypk1^19,31–33^. Yeast Orm1 and Orm2 are homologous on the primary sequence level but associated with different phenotypes. While the single deletion of *ORM1* has a minor effect on sphingolipid levels, deleting *ORM2* causes a more profound increase of LCBs^28,34^. Moreover, only Orm2 levels are regulated via the endosome and Golgi-associated degradation (EGAD) pathway, which depends on Ypk1-mediated phosphorylation of Orm2 and ER exit into the Golgi ^35^. The effects of Orm1 and Orm2 on sphingolipid levels and their different regulation suggest changes in their protein structures as a function of sequence divergence.

To identify the molecular mechanisms underlying Orm-mediated regulation of SPT activity and the functional distinctions between Orm1 and Orm2, we employ a combination of cryo-electron microscopy (cryo-EM) to resolve the structure of the SPOT-Orm2 complex, along with *in vivo* and *in vitro* assays. While the overall architecture and the ceramide-mediated inhibition of SPT activity in the SPOT-Orm2 complex is remarkably similar to the SPOT-Orm1 complex, we identify subtle differences in the binding of the two Orm proteins. Mutation of the Ypk1 phosphorylation sites of the Orm proteins affects their inhibitory function but does not alter the interaction of the Orm proteins with ceramides, suggesting that TORC2-Ypk1 signaling can still regulate SPT activity under ceramide-replete conditions. Our functional *in vitro* activity assays reveal lower SPT activity in the presence of Orm2 compared to Orm1. A newly developed assay for the inducible depletion of Orm proteins, combined with targeted lipidomics, yields similar effects *in vivo*. Together, our structural analysis of the SPOT-Orm2 complex, coupled with functional *in vivo* and *in vitro* assays, reveals a higher regulatory potential of the Orm2 protein towards SPT activity compared to Orm1.

## Results

For structure determination of the yeast SPT complex, including its negative regulator Orm2, we co-overexpressed Lcb1, Lcb2, Tsc3, Orm2, and Sac1 under the control of the inducible *GAL1* promoter in *Saccharomyces cerevisiae orm1*Δ *orm2*Δ cells (Fig. S 1A). A 3xFLAG epitope was inserted after P9 of Lcb1 to allow affinity purification while preserving its functionality^16^. Since S46, S47, and S48 in Orm2 are target sites for the yeast Ypk1 kinase^19,31^, and their phosphorylation is proposed to result in the dissociation of Orm2^23,35^, we mutated those residues to alanine to receive a non-phosphorylatable Orm2 mutant (Orm2^3A^). For cryo-EM studies, the SPOT-Orm2^3A^ complex was solubilized in glyco-diosgenin (GDN) and purified by FLAG-based affinity chromatography following our established protocol^21^ (Fig. S 1B). Mass spectrometric analysis confirmed the presence of all subunits with high sequence coverage (Fig. S 1C). Single particle cryo-EM revealed the compositional heterogeneity of the purified yeast SPT (Fig. S 6A, D). Within one dataset, we reconstructed a SPOT dimer (Fig. 1A, B, Fig. S6A) and a SPOT monomer complex (Fig. 1C, D, Fig. S6D). The monomeric SPOT-Orm2^3A^ was refined to a global resolution of 3.1 Å. By using C2-symmetry during non-uniform refinement, the SPOT-Orm2^3A^ dimer was resolved to 3.3 Å global resolution. Local refinement of a masked protomer further improved the resolution to 2.8 Å (Tab. S1, Fig. S 2, 3).

**Figure 1.**
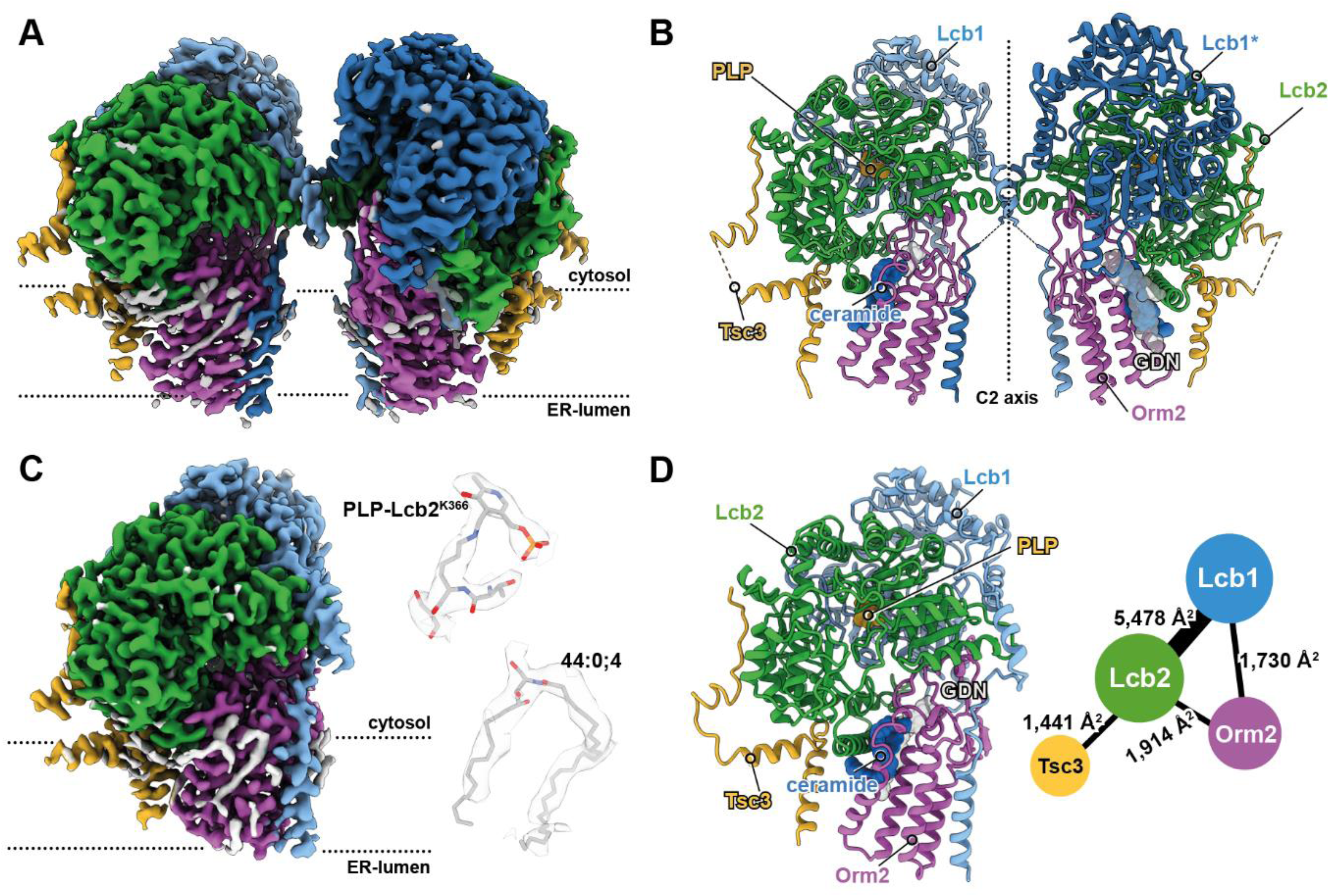
Architecture of yeast SPT-Orm2^3A^ complexes. **A**, Cryo-EM density map of C2 symmetric yeast SPOT-Orm2^3A^ dimer. **B**, Model of C2 symmetric yeast SPOT-Orm2^3A^ dimer at 3.3 Å resolution. **C**, Cryo-EM density map of yeast SPOT-Orm2^3A^ monomer and local densities of PLP bound to Lcb2-K366 and ceramide 44:0;4. **D**, Model of SPOT-Orm2^3A^ monomer at 3.1 Å resolution and a contact diagram of the subunits within the complex. The thickness of each line in the contact diagram is proportional to the contact area given in Å^2^ between subunits. All individual SPOT-Orm2^3A^ subunits are consistently colored: Lcb1 in blue, Lcb2 in green, Orm2 in purple, and Tsc3 in yellow. The membrane plane is indicated with dashed lines. Local densities are contoured at level 0.3.

### Structures of the SPOT-Orm2^3A^ monomer and dimer

The two catalytic subunits, Lcb1 and Lcb2, form the enzymatic core of the complex with a single transmembrane helix (TM1) of Lcb1 being resolved in the structure. Lcb2 is in contact with the membrane via an amphipathic helix. The Lcb2 lysine residue K366 is conserved across species and essential for SPT catalysis^36,37^. An apparent electron density for the PLP cofactor, covalently bound to the ε-amino group of Lcb2-K366 as an internal aldimine, was observed in the active site (Fig. 1B-D, Fig. S 4G, Fig. S 5A). The small regulatory subunit Tsc3 has a single transmembrane helix and exclusively interacts with Lcb2 via its elongated N-terminal region. An amphipathic helix of Tsc3 further increases the membrane contact of the complex. Lastly, Orm2^3A^ is positioned between the amphipathic helix of Lcb2 and the TM1 of Lcb1 but does not interact with Tsc3 (Fig. 1B, D). Analysis of the oligomeric states during data processing, showed that the distribution of monomeric versus dimeric species was about half and half for the SPOT-Orm2^3A^ complex (Fig. S 2). Although Sac1 was abundantly co-purified, a monomeric SPOT-Orm2^3A^ in complex with the PI4P phosphatase Sac1, as previously observed for SPOT-Orm1^3A^, could not be identified (Fig. S 2)^21^.

### Structural comparison of SPOT-Orm1^3A^ and SPOT-Orm2^3A^ complexes

Recent structural studies showed that the two transmembrane helices of the human SPTLC1 and *Arabidopsis thaliana At*Lcb1 subunits are swapped within dimers and establish a crossover^24–26^. However, our previously reported dimeric yeast SPOT-Orm1^3A^ complex does not include the Lcb1 transmembrane helices crossover^21^. To identify structural differences between Orm2 and its paralog Orm1, we compared our SPOT-Orm1^3A^ reconstructions with our new SPOT-Orm2^3A^ structures. In contrast to the dimeric SPOT-Orm1^3A^ complex, the dimeric SPOT-Orm2^3A^ did show a crossover of the Lcb1 transmembrane helices (Fig. 2B). Despite a loss of local density quality near the intra-membrane dimer interface, we could identify the crossover in our reconstruction by following the backbone-trace (Fig. S 4E). Additionally, analysis of the crossover region with OccuPy^38^ indicates a loss of local resolution due to mixed conformational occupancy (Fig. S 6C). Nevertheless, the presence of the Lcb1-crossover within the SPOT-Orm2^3A^ complex did not seem to foster dimerization, as the overall particle distribution with close-to-equal particle subsets of the dimer and monomer classes was similar to our previously resolved SPOT-Orm1^3A^ complex^21^. Besides the Lcb1 TM1 crossover, the overall architectures of the SPOT-Orm2^3A^ and the SPOT-Orm1^3A^ complexes are highly similar. Comparison of both dimeric complexes results in a global Cα root mean square (r.m.s.) deviation of only 0.41 Å (Fig. 2A), while comparison of both monomeric complexes has a Cα r.m.s. deviation of 0.77 Å (Fig. 2A).

**Figure 2.**
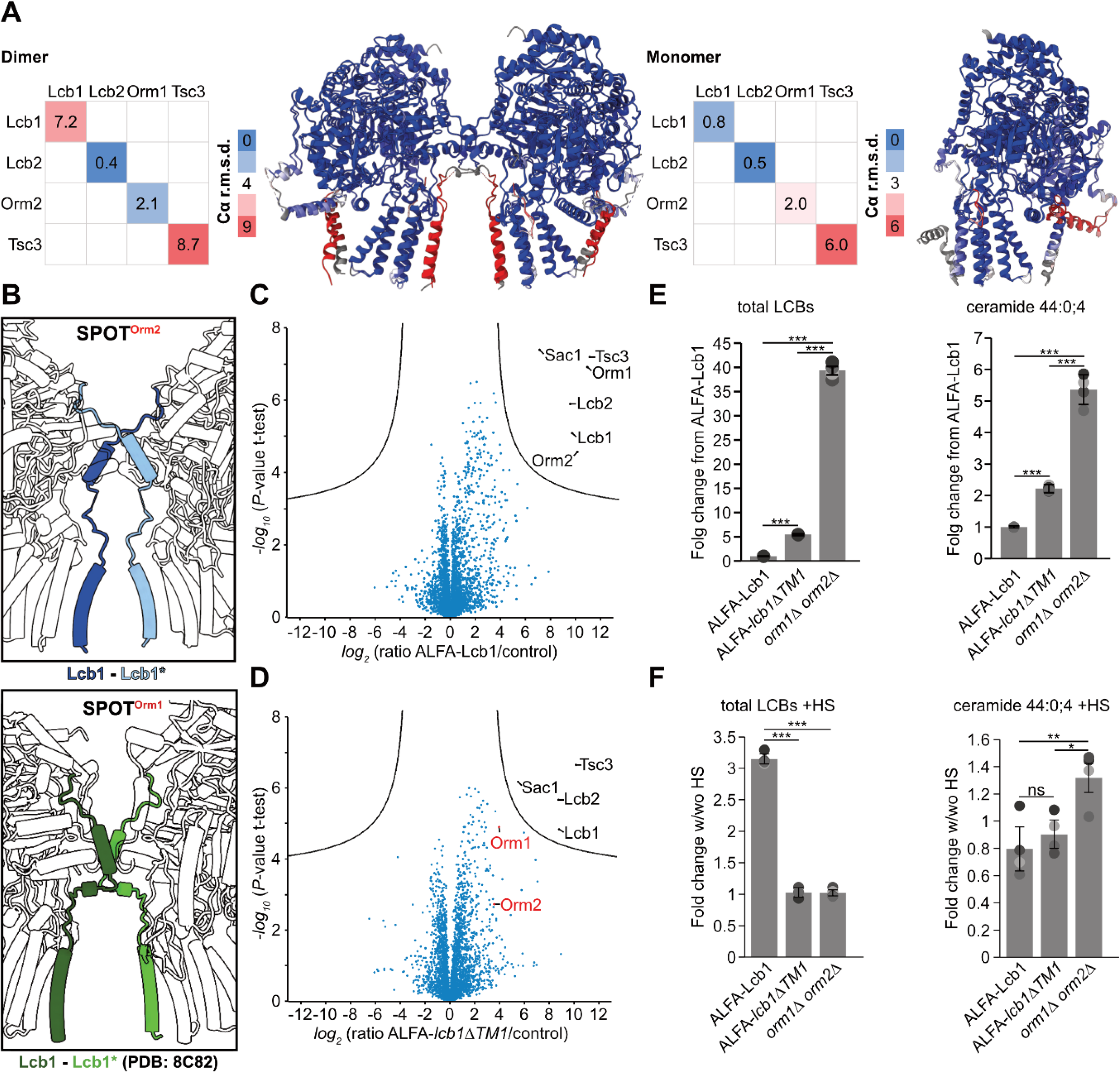
TM1 of Lcb1 is important for Orm binding to the SPT complex. **A**, Structural comparison of the dimeric and monomeric SPOT-Orm2 and SPOT-Orm1 (grey) complexes color-coded by Cα r.m.s deviation in Å. Local Cα r.m.s deviations in Å of single subunits are shown. Larger local differences with a high Cα r.m.s.d are shown in red. **B**, Close-up views of Lcb1-TM1 from SPOT-Orm2^3A^ dimer (Lcb1-TM1 in shades of blue) and SPOT-Orm1^3A^ dimer (Lcb1-TM1 in shades of grey; PDB: 8C82). Lcb1-TM1 from dimeric SPOT-Orm2^3A^ establishes a crossover, while Lcb1-TM1 from dimeric SPOT-Orm1^3A^ shows no crossover. **C**, Label-free proteomics of yeast cells expressing ALFA-Lcb1 compared to untagged control cells. In the volcano plot, the protein abundance ratios of ALFA-Lcb1 over control cells are plotted against the negative *log_10_* of the *P*-value of the two-tailed t-test for each protein. **D**, Label-free proteomics of yeast cells expressing ALFA-*lcb1ΔTM1* compared to untagged control cells. In the volcano plot, the protein abundance ratios of ALFA-*lcb1ΔTM1* over control cells are plotted against the negative *log_10_* of the *P*-value of the two-tailed t-test for each protein. **E**, Levels of total long-chain bases (LCBs) and ceramide 44:0;4 in *lcb1Δ* cells expressing ALFA-*LCB1* or ALFA-*lcb1ΔTM1* and *orm1Δ orm2Δ* cells. Displayed are the fold changes from ALFA-*LCB1*. Dots correspond to the values of four independent experiments. **F**, Levels of total long-chain bases (LCBs) and ceramide 44:0;4 in *lcb1Δ* cells expressing ALFA-*LCB1* or ALFA-*lcb1ΔTM1* and *orm1Δ orm2Δ* cells with and without (w/wo) 5 min of heat shock (HS) at 39°C. Displayed are the fold changes in LCBs and ceramide 44:0;4 levels after heat shock treatment over untreated controls. Dots correspond to the values of four independent experiments. Data in E and F were analyzed by one-way ANOVA with Tukey’s multiple-comparison test (*p < 0.05, **p < 0.01, ***p < 0.001). Exact *P*-values are shown in Tab. S4.

The interaction of the Orm proteins with Lcb1-TM1 is substantial for Orm-mediated regulation of SPT activity^7,23^. ALS-associated variants of SPTLC1 – the human homolog of yeast Lcb1 – are localized in TM1 and result in enhanced SPT activity due to an altered Orm-SPTLC1 interaction^7,8^. Additionally, deletion of amino acids H50-K85 in yeast Lcb1 disrupts its interaction with Orm proteins^23^. Based on our structural data, we generated a mutant lacking exactly TM1 (Y55-S73) of Lcb1 (*lcb1ΔTM1*). Using mass spectrometry-based proteomics, we first compared the interaction partners from ALFA-Lcb1 pulldowns and ALFA-*lcb1ΔTM1* pulldowns. While we significantly co-enriched all subunits in the WT pulldowns (Lcb2, Tsc3, Sac1, Orm1, Orm2; Fig. 2C), we observed a marked reduction of Orm1 and Orm2 binding in the pulldowns of the *lcb1ΔTM1* mutant (Fig. 2D). Moreover, Lcb1 mutant cells showed elevated levels of 3-KS and LCBs (Fig. 2E) and grew better in the presence of the SPT-specific inhibitor myriocin compared to control cells (Fig. S 1E). Both observations suggest a more active SPT in ALFA-*lcb1ΔTM1* cells, likely caused by the reduced binding of Orm proteins. However, Orm binding was not completely abolished in *lcb1ΔTM1* cells (Fig. 2D), which is also consistent with higher levels of 3-KS and LCBs and an increased resistance to myriocin observed for *orm1Δ orm2Δ* knockout cells than for *lcb1ΔTM1* cells (Fig. 2E, Sup-Fig. 1E). Our findings indicate that TM1 plays a crucial role in mediating the interaction between the SPT complex and Orm1/Orm2 but is not the only determinant for Orm binding. Indeed, the N-terminus of Orm2 shows additional interactions with both Lcb2 and Lcb1 (Fig. 1B, D). Moreover, Orm2 also interacts with the SPT via a ceramide bound between the amphipathic helix of Lcb2 and the transmembrane helices 1 and 2 of Orm2 (Fig. 1B, D). Notably, our analysis of the *lcb1ΔTM1* mutant also revealed higher levels of ceramides compared to WT cells. As Orm proteins are required for the ceramide-mediated inhibition of SPT activity, the elevated ceramide levels may well explain the reduced levels of all sphingolipid intermediates in the *lcb1ΔTM1* cells compared to the *orm1Δorm2Δ* cells (Fig. 2E). Accordingly, we also did not observe the previously shown elevation of sphingolipid intermediates after heat shock^34^ in the *lcb1ΔTM1* mutant compared to WT cells (Fig. 2F). This impaired response to heat shock in *lcb1ΔTM1* might also be accounted for higher basal levels of the inhibitory ceramides (Fig. 2E).

### Ceramide-binding is a conserved feature of SPT-regulation

Ceramides have been identified as common negative regulators of SPT activity from yeast to man^21,26,29^. Model building of SPOT-Orm2^3A^ revealed the presence of an elongated non-protein density within the monomeric and dimeric complex. Targeted lipidomics of the purified complex identified the long-chain ceramide 44:0;4, which was subsequently built into the cryo-EM densities (Fig. 1C, Fig. 3C, Fig. S 4F). Results from compositional heterogeneity analysis with OccuPy show an increase of mixed conformers within the periphery of the detergent-exposed portion of the ceramide acyl chains (Fig. S 6B, E). The ceramide is further stabilized through hydrophobic interactions with a structurally resolved GDN detergent molecule (Fig. S 4H, Fig. S 5C). The long-chain ceramide is positioned between the amphipathic helix (AH) of Lcb2 and TM1-2 of Orm2 (Fig. 3A, Fig. S 5B). The two conserved tyrosine residues of Lcb2 - Y485 close to the PATP-loop and Y110 – which we recently identified as gatekeeper residues, restrict the entry towards the active side near Lcb2-K366 in the SPOT-Orm2^3A^ complex, as previously shown for the SPOT-Orm1^3A^ complex^21^. Superposition of our previously resolved SPOT-Orm1^3A^ monomer with the SPOT-Orm2^3A^ monomer indicates a well-conserved binding mode of ceramides in the SPOT-complex (Fig. 3B). Homologs from human and *Arabidopsis thaliana* SPTs in complex with ceramides share this general binding mode^26,29,30^, indicating a conserved inhibitory function. Interestingly, the ceramide is located near the N-terminal loop, which includes the Ypk1 phosphorylation target sites (Fig. S 7C). To test whether the differences in SPT inhibition of Orm2 mutants^35,39^, whose three Ypk1 targeted serines (S46-48) are mutated to either alanines (Orm2^3A^) or phospho-mimetic aspartates (Orm2^3D^), resulted from distinct levels of bound ceramide, we purified the SPOTS-Orm2^3A^ and SPOTS-Orm2^3D^ complexes. Similar to the SPOTS-Orm2^3A^ complex, the SPOTS-Orm2^3D^ complex was stable after purification in GDN (Fig. S 1F). The purified complexes were subjected to lipid extraction and targeted lipidomics. As a control, the SPOTS-Orm1^3A^ complex was used. No differences in the typical yeast ceramide 44:0;4 were observed between the Orm2^3A^ and Orm2^3D^ complexes (Fig. 3C), indicating that Ypk1-mediated phosphorylation has a different effect than regulating bound ceramide levels.

**Figure 3.**
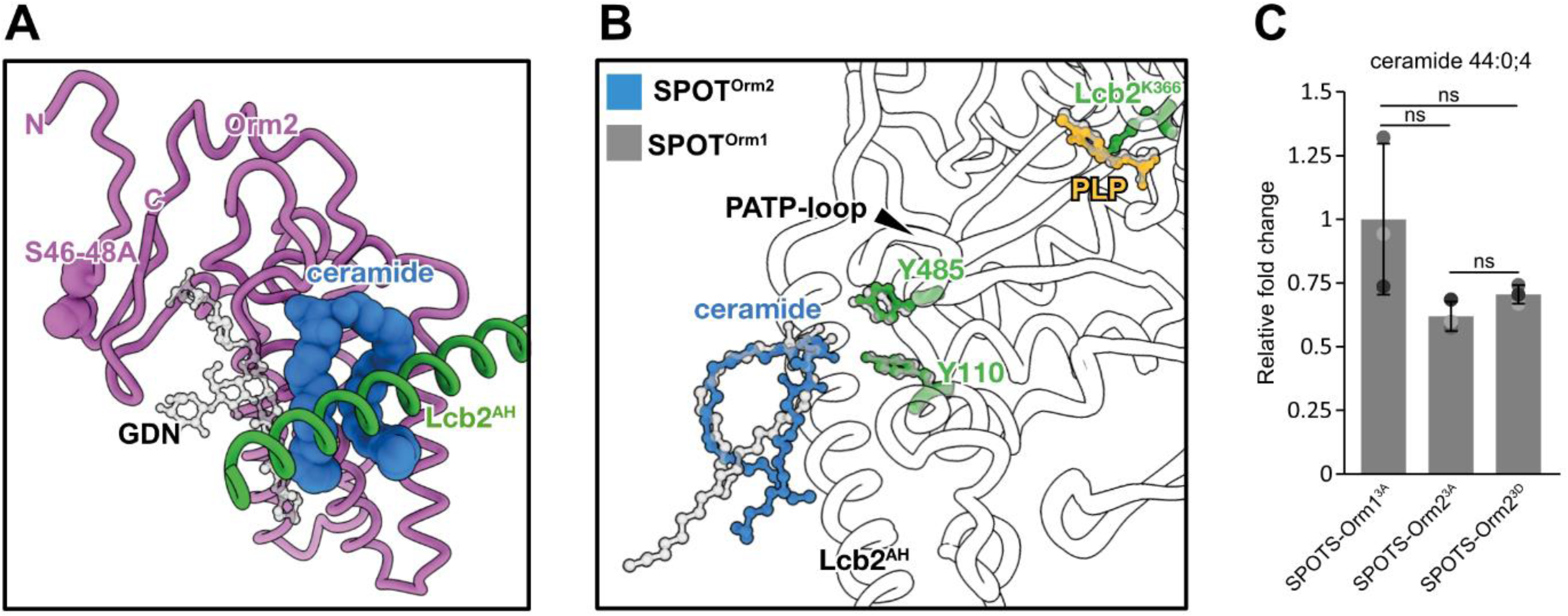
Ceramide-binding mode in yeast SPOT complexes. **A**, Binding mode of ceramide (blue) sandwiched between Orm2 (purple) and the amphipathic helix (AH) of Lcb2 (green). The phosphorylation sites S46-48 mutated to alanine are shown as spheres. The bound ceramide 44:0;4 is further stabilized by a GDN molecule (transparent). **B**, Superposition of yeast SPOT complexes. Ceramide 44:0;4 is shown in blue, and the gatekeeper residues Y485 and Y110 (green) and the internal aldimine of Lcb2-K366 (green) and PLP (yellow) are shown as ball-and sticks. The SPOT-Orm1 complex is shown in transparent. (SPOT-Orm1: PDB 8C80). **C**, Ceramide 44:0;4 quantification from GDN-solubilized and purified SPT complexes shown as relative fold change to SPOTS-Orm1^3A^. n=3 technically independent samples and data are presented as mean values ± SD. ns, not significant. Data were analyzed by one-way ANOVA with Tukey’s multiple-comparison test (ns > 0.05). Exact *P*-values are shown in Tab. S4

### Orm1 and Orm2 have distinct inhibitory potentials on SPT activity *in vitro*

The primary sequence of yeast Orm1 and Orm2 is highly similar, with 67 % sequence identity and 78 % sequence similarity (Matrix: EBLOSUM62) (Fig. 4A). When comparing the overall composition of both SPT complexes with Orm1 or Orm2, the two Orm proteins within the structures show a Cα r.m.s deviation of only 0.7 Å (Fig. 4B). However, larger local structural differences can be observed within the C-terminal regions (Fig. 4C) and within a short α-helix, that interacts with Lcb1 and Lcb2 (Fig. 4D). Notably, the first 36 amino acids of Orm2 (Fig. 4A) and the first 38 amino acids of Orm1, which harbor the strongest divergence in primary sequence between both Orm proteins, could not be resolved in our structures^21^ (Fig. S 7A, B). Therefore, further structural differences in the N-terminal parts are possible, and previous studies have already hinted at an important role for the N-terminus in the inhibitory function of Orm proteins in other species^26,29^. To investigate differences in the Orm1- and Orm2-containing complexes, we analyzed the activities of the purified complexes using *in vitro* activity assays. Interestingly, the inhibitory effect of Orm1 on SPT activity appears to be lower than that of Orm2 (Fig. 4E). The purified SPOTS-Orm2^3A^ complex showed a nearly 7-fold lower SPT activity than the SPOTS-Orm1^3A^ complex *in vitro,* and both complexes were sensitive to myriocin (Fig. 4E). Therefore, the observed differences in SPT activities suggest a divergence of function, resulting from the apparent sequence or conformational differences of yeast Orm proteins. The SPOTS-Orm2^3D^ complex showed a 7-fold increase in enzymatic activity compared to the purified SPOTS-Orm2^3A^ complex. Its activity was likewise abolished upon myriocin treatment (Fig. 4E). While previous *in vivo* experiments (see also Figure 5E) have observed dissociation of Orm proteins upon phosphorylation^23,35^, our findings indicate that the phospho-mimetic Orm2 mutant can remain associated with the SPT complex apparently without compromising its activation.

**Figure 4.**
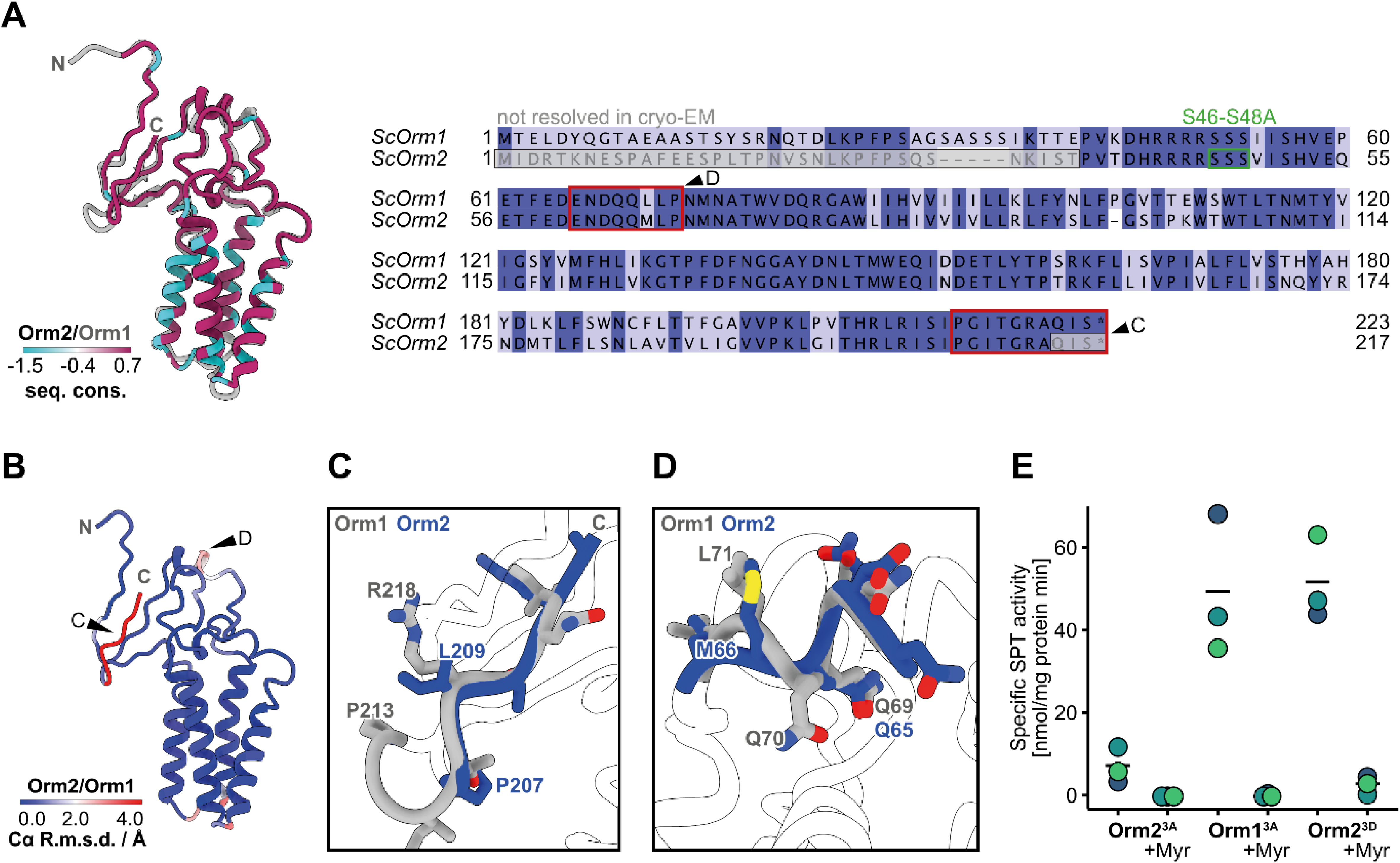
Orm proteins are highly homologues but show functional divergence. **A**, Cartoon representation of yeast Orm1 and Orm2 color-coded by sequence conservation and pair-wise sequence alignment of *S. cerevisiae* Orm1 (P53224) and Orm2 (Q06144) color-coded by sequence identity. The mutated phosphorylation sites S46-48A are marked in green. Non-resolvable residues from cryo-EM are marked in light gray. Sequences are color-coded by conservation with a cut-off at 30 %. Sequences in red correspond to regions of high Cα r.m.s.d. Underlying multiple sequence alignments were prepared with Jalview^40^. **B**, Cartoon representation of yeast Orm1 and Orm2 color-coded by Cα r.m.s deviation in Å. **C, D** Regions of high Cα r.m.s.d. within the C-terminal sequence (C) and a central short α-helix (D) of aligned Orm1 and Orm2 structures. **E**, Specific SPT enzyme activity measurement of SPOTS complexes containing the indicated Orm2 and Orm1 variants with or without the specific SPT-inhibitor myriocin (Myr). Data is presented as mean values and dots correspond to values of three technically independent samples (n=3).

**Figure 5.**
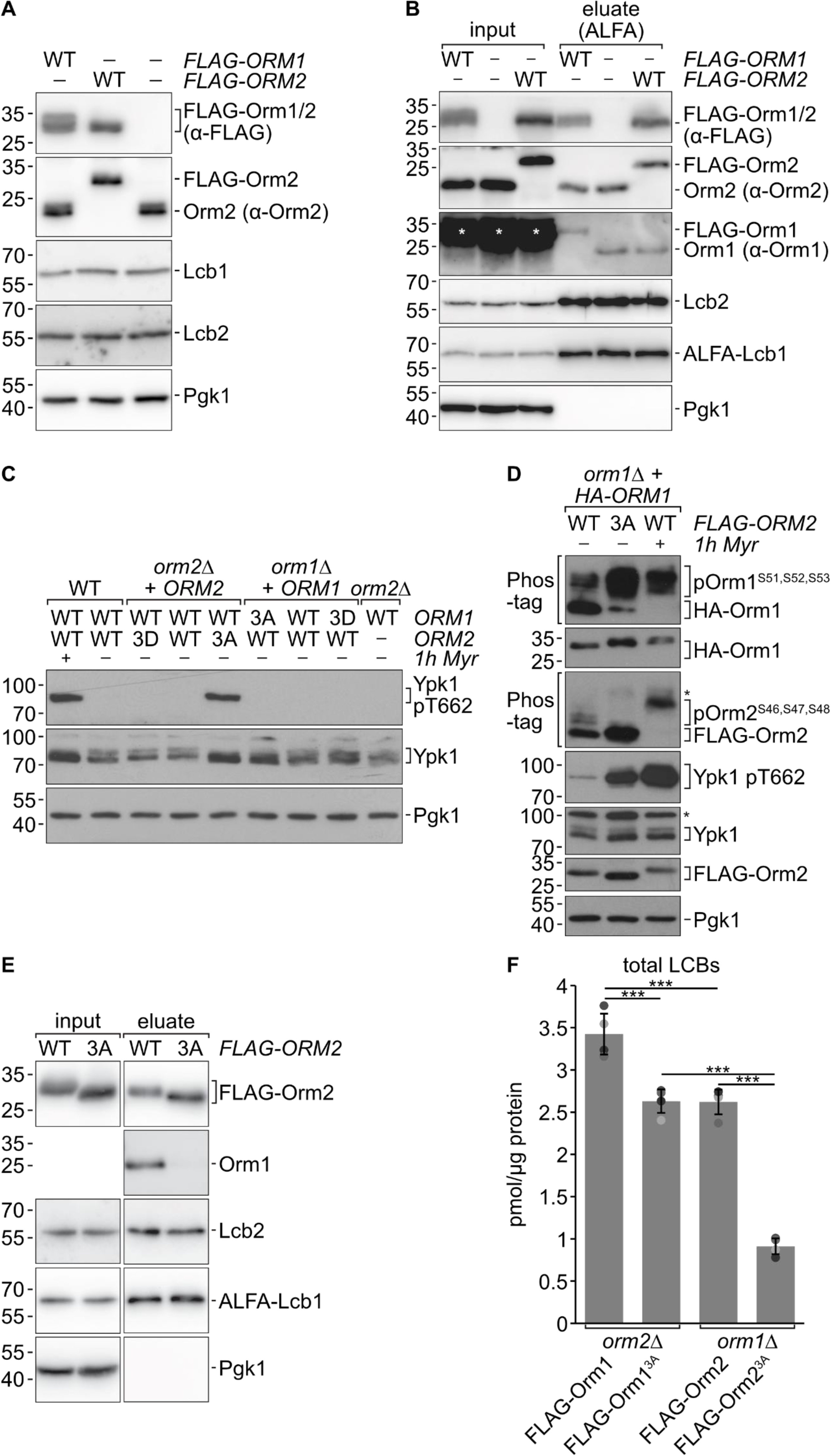
Yeast Orm1 and Orm2 exhibit different inhibitory potentials towards SPT activity. **A**, Whole-cell protein extracts of WT cells and cells chromosomally expressing *FLAG-ORM1* or *FLAG-ORM2* from their endogenous loci analyzed by SDS PAGE and Western blotting with the indicated antibodies. **B**, Input and eluate fractions of a native immunoprecipitation of ALFA-Lcb1 from WT cells and cells chromosomally expressing *FLAG-ORM1* or *FLAG-ORM2* from their endogenous loci analyzed by SDS PAGE and Western blotting with the indicated antibodies. * indicates a cross-reaction of the Orm1 antiserum. **C**, Whole cell protein extracts of WT cells or *orm1*Δ and *orm2*Δ cells harboring centromeric plasmids of the indicated *ORM1* or *ORM2* variants (expressed from their endogenous promoter and terminator) analyzed as in A. Where indicated, cells were treated with 2.5 µM myriocin (in DMSO) for one hour. **D**, Whole cell protein extracts of *orm1*Δ cells expressing *3xHA-ORM1* from a centromeric plasmid and chromosomally expressing *FLAG-ORM2* WT or 3A mutant from the endogenous locus analyzed as in C. * denotes cross-reactive bands. **E**, Input and eluate fractions of a native immunoprecipitation of ALFA-Lcb1 from cells chromosomally expressing *FLAG-ORM2* WT or the 3A mutant from the endogenous locus analyzed by SDS PAGE and Western blotting with the indicated antibodies. **F**, Levels of total long-chain bases (LCBs) in *orm2*Δ cells expressing *FLAG-ORM1* WT or *FLAG-ORM1*^3A^ and *orm1*Δ cells expressing *FLAG-ORM2* WT or *FLAG-ORM2*^3A^. mean ± s.d. (n = 4). Data were analyzed by one-way ANOVA with Tukey’s multiple-comparison test (*p < 0.05, **p < 0.01, ***p < 0.001). Exact *P*-values are shown in Tab. S4

Interestingly, the N-terminus of Orm2 containing the activating phosphorylation sites is located at a surface of the SPT complex, which already has a negative electrostatic surface potential (Fig. S 7D). Additional phosphorylation of the Ypk1 target sites might lead to conformational changes of the Orm2 N-terminus due to repulsive forces. Notably, the short α-helix of Orm2 at the interface with Lcb1 and Lcb2 that we observed in a different conformation between our more active SPOT-Orm1^3A^ and less active SPOT-Orm2^3A^ complexes is also closely adjunct (Fig. 4D, Fig. S 7E). Hence, it is tempting to speculate that phosphorylation-induced changes in this helix might underly the activation of SPT activity.

### Orm1 and Orm2 differentially regulate SPT activity *in vivo*

Poised by the different regulatory effects of Orm1 and Orm2 *in vitro,* we investigated their regulation of the SPT *in vivo*. Regulatory effects have been attributed to variations in expression in the past^39^. Therefore, we initially analyzed the expression levels of Orm1 and Orm2. N-terminally 3xFLAG-tagged Orm1 and Orm2 expressed from their respective endogenous genomic loci exhibited comparable protein levels in total cell extracts (Fig. 5A). Alternatively, differences in the quantities of bound Orm1 and Orm2 could lead to changes in SPT activity. To test this, we analyzed the levels of co-purified Orm1 and Orm2 in Lcb1 pulldowns. Purification of the SPOTS complex via ALFA-tagged Lcb1 co-purified similar amounts of FLAG-Orm1 and FLAG-Orm2 (Fig. 5B). We have previously suggested that the SPOTS monomer is the prevalent form of the complex *in vivo*^21^, implying that SPOTS-Orm1 and SPOTS-Orm2 complexes were also similarly abundant in cells. Moreover, the levels of co-purified FLAG-Orm1 and FLAG-Orm2 were similar to the amounts of the co-purified, untagged Orm proteins when the eluates were analyzed with specific antisera, suggesting that the FLAG-tagged variants accurately reported the native binding levels of the two Orm proteins to the SPT. Since the expression levels of both Orm proteins and the abundance of the SPOTS complexes were similar, we analyzed their inhibitory effect on SPT activity *in vivo*. We first analyzed the effects of the Orm phospho-site-mutants on the homeostatic TORC2-Ypk1 signaling axis. Expression of Orm2^3A^ resulted in strong phosphorylation of the activating TORC2 phosphorylation site of Ypk1 (T662), as observed previously^35^ (Fig. 5C). The activation of Ypk1 was comparable to the treatment with myriocin, suggesting that the expression of Orm2^3A^ dampened the synthesis of sphingolipids in cells sufficiently to activate a feedback response via TORC2. In contrast, expression of the Orm1^3A^ mutant did not increase the levels of Ypk1 phosphorylation. The Orm2^3A^ mutant exhibits a higher affinity for SPT *in vivo compared to Orm2 WT* ^23,35^, and a similar trend was noted for Orm1^3A^ (Fig. S 8A). Hence, lower activation of TORC2-Ypk1 signaling is not due to a failure of unphosphorylated Orm1 to engage the SPT more strongly.

When sphingolipid levels are low, Ypk1 phosphorylates the regulatory Orm proteins to increase LCB biosynthesis and restore sphingolipid levels^19,31^. This regulation is accompanied by a reduced binding of the Orm proteins to the SPT complex, which was proposed to release SPT inhibition^23,35^. Indeed, expression of Orm2^3A^ and the ensuing hyperactivation of Ypk1 caused hyper-phosphorylation of Orm1 (Fig. 5D), efficiently depleting Orm1 from the SPOTS complexes (Fig. 5E). Thus, when cells express Orm2^3A^, they react by adjusting TORC2-Ypk1 signaling to decrease the amount of Orm1 within SPOTS complexes and thereby presumably boost sphingolipid biosynthesis, whereas Orm1^3A^ did not provoke a similar response (Fig. S 8B). This indicated differences in the steady state activity of Orm1^3A^- and Orm2^3A^-containing SPOTS complexes, as our *in vitro* analysis suggested. Yet, steady-state levels of LCBs in cells expressing Orm1^3A^ or Orm2^3A^ were previously found to be roughly similar^34^, presumably due to the here-observed compensatory effects via TORC2-Ypk1 signaling. Therefore, we aimed to quantify sphingolipid levels in cells expressing Orm1^3A^ and Orm2^3A^ without interfering with these confounding effects by deleting the corresponding *ORM* gene. Simultaneous mutations of *ORM1* and *ORM2* are, however, often rapidly suppressed. The substantial growth defect of *orm1Δ orm2Δ* double mutants is typically ameliorated within a few generations. We, therefore, devised an inducible gene depletion strategy (Fig. S 8C). We generated diploid yeast strains that were heterozygous for the WT or mutants of *ORM1* or *ORM2*, as well as for the deletion of the respective *ORM1* or *ORM2* gene. Before generating the haploid double mutants, we transformed these diploids with a cover plasmid expressing *ORM1*-WT alongside a counter-selectable *URA3* marker.

These strains were subsequently used to generate cells that expressed only a single version of the SPOTS complex, either containing Orm1 WT, Orm2 WT, or their respective 3A or 3D mutants, by forcing loss of the WT cover plasmid on a medium containing 5-fluoroacetic acid (5-FOA). While all strains grew equally well in the presence of the WT cover plasmid (Fig. S 9A), an induced *orm1Δ orm2Δ* double deletion strain showed the expected severe growth defect on 5-FOA medium (Fig. S 9B). A similar growth defect was observed for cells expressing only the phospho-mimetic 3D mutants of Orm1 or Orm2 (Fig. S 9B), confirming that these mutants could no longer restrain SPT activity, similar to the deletion of both *ORM* genes. Notably, after 3-6 days, suppressor clones that had overcome growth inhibition reproducibly appeared on the plates, while a control strain deleted for the essential *LCB1* gene and rescued with a corresponding cover plasmid never showed suppression (Fig. S 9C). In contrast, cells containing *ORM1* WT, *ORM1*^3A^, *ORM2* WT, or *ORM2^3A^* as the only *ORM* gene copy supported growth on 5-FOA (Fig. S 9B). We confirmed that cells grown for 48 hours in the presence of 5-FOA (i.e., before the typical onset of phenotypic suppression) had lost detectable expression of untagged Orm1 in Western blot (Fig. S 9D).

We measured steady-state LCB levels of cells containing Orm1^3A^ and Orm2^3A^ as the sole Orm protein (Fig. 5 F). As expected, cells expressing only Orm1 WT (i.e., *orm2Δ* cells) had mildly higher LCB levels than those expressing only Orm2 WT (i.e., *orm1Δ* cells). Expression of only Orm2^3A^ caused a severe decrease of LCB levels (both DHS and PHS species) to approximately 35% of the corresponding WT levels. In contrast, cells with Orm1^3A^ expressed as the only Orm protein now showed a clear difference, as their LCB levels were only approximately 75% of the Orm1 WT control. Our genetic setup for the induced generation of cells expressing only one Orm variant revealed that Orm1 and Orm2 have differences in their capacity to inhibit SPT activity. While unphosphorylated Orm2 strongly repressed SPT activity, we observed a lower potential of unphosphorylated Orm1 to inhibit SPT activity *in vivo*, consistent with our *in vitro* analysis of the isolated complexes (Fig. 4E). Therefore, Orm2 is a more potent inhibitor of SPT activity compared to Orm1.

## Discussion

The Orm proteins are evolutionarily conserved regulators of the SPT and are essential to maintaining sphingolipid homeostasis. However, their regulation appears to differ between species and the highly homologous yeast proteins Orm1 and Orm2. Here, we present the structure of the yeast SPOT complex harboring the non-phosphorylatable Orm2^3A^ mutant in both monomeric and dimeric states. We use this structure to compare the overall architecture of the Orm2^3A^-containing complex to the previously solved structures of the Orm1^3A^-containing SPOT complexes. While the complexes are generally very similar, Orm2 and Orm1 show different regulatory capacities of the SPT *in vivo* and *in vitro*.

A notable distinction between the Orm2-containing complex and the Orm1-containing complex is the absence of the PI4P phosphatase Sac1 in the structure of the Orm2 complex. While Sac1 was initially detectable in all purifications, we did not observe any particles containing Sac1. Whether the absence of Sac1 has a biological relevance is unclear. It is also plausible that Sac1 is lost during grid preparation, possibly linked to its relatively small interaction interface within the complex^21^. Furthermore, the dimeric Orm2-containing complex shows a helix swap of the Lcb1-TM1, similar to observations in human and *A. thaliana* SPT-Orm complexes^24–26^ and not for the Orm1-containing complex^21^. However, our previous findings demonstrated that the predominant form of the SPOTS complex *in vivo* is its monomeric state^21^. Therefore, the biological significance of the helix swap in the dimeric complex is unclear and might also be attributed to the strong overexpression of the complexes for cryo-EM sample preparation. In support of this argument, mutation of the interface does not change the activity of the complex *in vitro*^24^. Additionally, we could not resolve the N-terminal TM0 helix of Lcb1 in the monomeric SPOT-Orm2 complex that we observed in the monomeric Orm1-containing complex^21^. The N-terminus of Lcb1 was shown to interact with several ergosterol molecules within the SPOTS-Orm1 complex. The functional relevance of this interaction is unclear, although a cross-regulation of sphingolipids and sterols has been suggested previously^41,42^. Whether the SPOT-Orm2 complex also binds ergosterol or whether the loss of resolution in this region in the SPOT-Orm2 complex has a biological cause remains to be determined.

The TM1 of Lcb1 is essential for the proper ceramide- and Orm-mediated regulation of the SPT, and mutations in the TM1 of human SPTLC1 are associated with early onset ALS^7,8^. However, even a full deletion of the Lcb1-TM1 did not completely abolish the binding of the Orm proteins, suggesting that interactions with other parts of Lcb1 and Lcb2 are sufficient to maintain a fraction of Orm proteins in their position in the complex. Accordingly, deletion of just TM1 (residues 55-73) of Lcb1 did not phenocopy an *orm1Δ orm2Δ* double deletion in our hands, in contrast to Han et al., 2019, who deleted a larger part of Lcb1 (residues 50-85) that did no longer support Orm binding^23^. Our data is consistent with work in human cells, suggesting that ALS mutations in the TM1 of SPTLC1 also did not entirely abrogate the binding of ORMDL proteins to the SPT and its ORMDL-mediated regulation^8^. Moreover, we noted increased ceramide levels in cells with deleted TM1 of Lcb1, which may additionally help to stabilize Orm proteins at the SPT in this setting.

While the overall architecture of the Orm2-containing SPOT complex is very comparable to the Orm1-containing complex, we can now show for the first time a clear, distinct function of Orm1 and Orm2 in regulating SPT activity. Orm2 is a more potent inhibitor of SPT activity. Orm2^3A^ bound to the SPOTS complex demonstrates significantly stronger inhibition of SPT *in vivo* and *in vitro*. We observed similar effects before^34^. However, we can separate the Orm1 and Orm2 function *in vivo* with our newly developed plasmid covering system. Our data support a model in which Ypk1 phosphorylation of the Orm proteins modifies their interaction with the SPT. This modification leads to either the release of phosphorylated Orm proteins from the SPT, a reduction in interaction, or both. Notably, even the phospho-mimetic Orm2^3D^ protein can still interact with the SPT, as evidenced by the purification of a stable Orm2^3D^-containing SPOTS complex. Notably, the SPOTS-Orm2^3D^ complex exhibits significantly higher activity than the SPOTS-Orm2^3A^ complex. Thus, phosphorylation by Ypk1 may induce a conformational change translating into a reorientation of a short α-helix of Orm2, which interacts with Lcb1 and Lcb2 and shows profound local structural differences to Orm1 (Fig. 4D, Fig. S7E). Possibly, in the SPOT-Orm1^3A^ complex, the conformation of this α-helix is more favorable for allowing SPT catalysis than in the SPOT-Orm2^3A^ complex, explaining why the former retains significant activity while the latter is almost inactive. Hence, the release of Orm proteins from the SPT may not be required to exert the acute stimulatory effect of Ypk1 phosphorylation on SPT activity, which conformational rearrangements can already achieve. However, the release of Orm2 from the SPOTS complex, or preventing the interaction of phosphorylated Orm2 in the first instance, is the prerequisite for its ER export and turnover by the endosome and Golgi-associated degradation. The EGAD pathway thus might serve as a mechanism to down-regulate Orm2 protein levels and adjust sphingolipid homeostasis on a longer timescale^35^.

Interestingly, mimicking phosphorylation of Orm2 also did not lead to a loss of ceramide from the SPOTS complex, suggesting that these are independent regulatory mechanisms that both converge on SPT activity. While ceramide-mediated SPT inhibition via Orm proteins is an evolutionary conserved process^21,24–26^, its mechanistic details and especially its regulation remain unclear. A physiological condition that leads to increased SPT activity in yeast is heat exposure^34,43^. We can only speculate that, in addition to the documented increase in Ypk1 phosphorylation under these conditions, which can only partially account for the increased SPT activity^32^, the elevated temperature might result in increased membrane fluidity^44^, facilitating the dissociation of phosphorylated Orms and ceramide.

Whether the phosphorylation of the Orm proteins leads to a conformational change in the proteins that changes their affinity for the complex can only be finally resolved in structural studies of the phosphorylated or phospho-mimetic Orm proteins. However, we can now clearly show distinct roles for the two yeast Orm proteins in regulating SPT activity. Similarly, we speculate that the human orthologues ORMDL1-3 will only have partially overlapping functions since the respective ORMDL1-3 deficient cells and mouse models also show distinct phenotypes^45,46^. Hence, it will be interesting to see if the three different ORMDL isoforms in mammalian cells also have different capacities for SPT inhibition.

## STAR Methods

### Key resources table

| REAGENT or RESOURCE | SOURCE | IDENTIFIER |
| --- | --- | --- |
| **Antibodies** |  |  |
| Goat anti-mouse IgG-Peroxidase | Sigma | Cat. # A4416;<br>RRID:AB_258167 |
| Goat anti-rabbit IgG-Peroxidase | Sigma | Cat. # A0545;<br>RRID:AB_257896 |
| Rabbit anti-goat IgG-Peroxidase | Sigma | Cat. # A5420 |
| Mouse monoclonal anti PGK1 (22C5D8) | Invitrogen | Cat. # 459250;<br>RRID:AB_2532235 |
| Mouse monoclonal anti-FLAG M2 | Sigma | Cat. # F3165;<br>RRID:AB_259529 |
| Mouse monoclonal anti-HA (IgG1K, clone 16B12) | Bio Legend | Cat. # 901502;<br>RRID:AB_2565006 |
| Mouse monoclonal anti-Ypk1 phospho-T662 | Gift from Loewith lab | (Berchtold et al., 2012)<sup>32</sup> |
| Goat polyclonal anti-Ypk1 | Santa Cruz | Cat. # yl15;<br>RRID:AB_793265 |
| Rabbit polyclonal anti-ALFA | NanoTag<br>Biotechnologies | Cat. #N1581 |
| Rabbit polyclonal anti-Lcb1 | Gift from Teresa Dunn | (Gable et al., 2000)<sup>16</sup> |
| Rabbit polyclonal anti-Lcb2 | Gift from Teresa Dunn | (Gable et al., 2000)<sup>16</sup> |
| Rabbit polyclonal anti-Orm1 purified | Gift from Teresa Dunn | (Han et al., 2019)<sup>23</sup> |
| Rabbit polyclonal anti-Orm2 purified | Gift from Teresa Dunn | (Han et al., 2019)<sup>23</sup> |
| **Chemicals, peptides, and recombinant proteins** |  |  |
| Myriocin | Sigma-Aldrich | M1177-5MG |
| Glyco-diosgenin (GDN) | Anatrace | GDN101 5 gm |
| Phenylmethylsulfonyl fluoride (PMSF) | Roth | 6367.2 |
| FLAG M2 agarose beads | Sigma-Aldrich | A2220-25ml |
| 3xFLAG peptide | Biozol | A6001 |
| Palmitoyl coenzyme A lithium salt | Sigma-Aldrich | P9716 |
| Pyridoxal 5'-phosphate hydrate | Sigma-Aldrich | P9255 |
| L-serine | Merck KGaA | 56-45-1 |
| 5-Fluoroacetic acid | Thermo Fisher | Cat. # R0812 |
| anti-ALFA selector ST magnetic beads | NanoTag<br>Biotechnologies | Cat. # N1516 |
| glyco-diosgenin (GDN) | Sigma | Cat. # 850525 |
| 3xFLAG elution peptide | Sigma | Cat. # F4799 |
| Phos-tag acrylamide | Wako | Cat. # 304-93521 |
| PhosStop | Roche | Cat. # 04906845001 |
| **Critical commercial assays** |  |  |
| Coenzyme-A(CoA)-Assay-Kit | Sigma-Aldrich | MAK034 |
| **Deposited data** |  |  |
| ScSPT-Orm2 dimer; model | This study | PDB-ID 8QOF |
| ScSPT-Orm2 dimer; map | This study | EMD-18536 |
| ScSPT-Orm2 dimer; masked map | This study | EMD-18533 |
| ScSPT-Orm2 monomer; model | This study | PDB-ID 8QOG |
| ScSPT-Orm2 monomer; map | This study | EMD-18537 |
| Mass-Spectrometry Data of ALFA-Lcb1 and ALFA-*lcb1*ΔTM pulldowns | This study | PXD048754 |
| **Experimental models: Organisms/strains** |  |  |
| MATa/MATα leu2-3,112 trp1-1 can1-100 ura3-1 ade2-1 his3-11,15 | Walther et al., 2007<sup>47</sup> | W303 |
| MATa leu2-3,112 ura3-52 his3-Δ200 trp1-Δ901 suc2-Δ9 lys2-801 | Robinson et al., 1988<sup>48</sup> | SEY6210.1 |
| W303 MATa/MATα *orm1Δ::natNT2* *orm2Δ::hphNT1* pRS404-GAL1-TSC3::TRP1 pRS406-GAL1-ORM1-3A::URA3 pRS406-GAL1-3xFLAG-LCB1::URA3 pRS405-GAL1-LCB2::LEU2 pRS403-GAL1-SAC1::HIS3 | Schäfer et al., 2023<sup>21</sup> | FFY4192 |
| W303 MATa/MATα *orm1Δ::natNT2* *orm2Δ::hphNT1* pRS404-GAL1-TSC3::TRP1 pRS405-GAL1-ORM2-3A::LEU2 pRS406-GAL1-3xFLAG-LCB1::URA3 pRS405-GAL1-LCB2::LEU2 pRS403-GAL1-SAC1::HIS3 | This study | FFY4193 |
| W303 MATa/MATα *orm1Δ::natNT2* *orm2Δ::hphNT1* pRS404-GAL1-TSC3::TRP1 pRS405-GAL1-ORM2-3D::LEU2 pRS406-GAL1-3xFLAG-LCB1::URA3 pRS405-GAL1-LCB2::LEU2 pRS403-GAL1-SAC1::HIS3 | This study | FFY5280 |
| SEY6210 MATα pRS406::URA3 | This study | FFY4631 |
| SEY6210 MATα *lcb1Δ::hphNT1* pRS406-ALFA-Lcb1::URA3 | This study | FFY3777 |
| SEY6210 MATα ade2-101 *lcb1Δ::hphNT1* pRS406-ALFA-lcb1DTM | This study | FFY4540 |
| SEY6210 MATα leu2-3,112 ura3-52 his3-Δ200 trp1-Δ901 suc2-Δ9 lys2-801; GAL | Robinson et al., 1988<sup>48</sup> | SEY6210 |
| SEY6210 MATα LYS2 *orm1Δ::natNT2* *orm2Δ::kanMX6* | Esch et al., 2023<sup>34</sup> | FFY2354 |
| SEY6210.1 MATa *orm1::3x-FLAG-ORM1* (TRP1) | Esch et al., 2023<sup>34</sup> | OSY928 |
| SEY6210.1 MATa *orm2::3x-FLAG-ORM2* (TRP1) | Esch et al., 2023<sup>34</sup> | TMY002 |
| SEY6210.1 MATa lcb1::TRP1 + pVN003 (ALFA-LCB1/pRS415) | This study | OSY1048 |
| SEY6210.1 MATa lcb1::TRP1 orm1::3x-FLAG-ORM1 (TRP1) + pVN003 (ALFA-LCB1/pRS415) | This study | OSY1052 |
| SEY6210.1 MATa lcb1::TRP1 orm2::3x-FLAG-ORM2 (TRP1) + pVN003 (ALFA-LCB1/pRS415) | This study | OSY1074 |
| SEY6210.1 MATa lcb1::TRP1 orm1::3x-FLAG-ORM1-3A (TRP1) + pVN003 (ALFA-LCB1/pRS415) | This study | OSY1091 |
| SEY6210.1 MATa lcb1::TRP1 orm2::3x-FLAG-ORM2-3A (TRP1) + pVN003 (ALFA-LCB1/pRS415) | This study | OSY1080 |
| SEY6210.1 MATa orm1::3x-FLAG-ORM1-3A (TRP1) | This study | VNY005 |
| SEY6210.1 MATa orm1::HIS3 | Schmidt et al., 2019<sup>35</sup> | OSY510 |
| SEY6210.1 MATa orm1::TRP1 | Schmidt et al., 2019<sup>35</sup> | OSY774 |
| SEY6210.1 MATa orm1::HIS3, orm2::3x-FLAG-ORM2-3A (TRP1) | This study | TMY084 |
| SEY6210.1 MATa orm1::HIS3, orm2::3x-FLAG-ORM2-3A (TRP1) | This study | OSY912 |
| SEY6210.1 MATa orm1::HIS3, orm2::3xFLAG-ORM2-WT (TRP1) + ORM1/416 (pVB23) | This study | OSY1058 |
| SEY6210.1 MATa orm1::HIS3, orm2::3xFLAG-ORM2-3A (TRP1) + ORM1/416 (pVB23) | This study | OSY1060 |
| SEY6210.1 MATa orm1::HIS3, orm2::3xFLAG-ORM2-3D (TRP1) + ORM1/416 (pVB23) | This study | OSY1062 |
| SEY6210.1 MATa orm1::HIS3, orm2::TRP1 + ORM1/416 (pVB23) | This study | OSY1064 |
| SEY6210.1 MATa orm1::3xFLAG-ORM1-WT (TRP1), orm2::TRP1 + pRS413 + ORM1/416 (pVB23) | This study | OSY1161 |
| SEY6210.1 MATa orm1::3xFLAG-ORM1-3A (TRP1), orm2::TRP1 + pRS413 + ORM1/416 (pVB23) | This study | OSY1162 |
| SEY6210.1 MATa orm1::3xFLAG-ORM1-3D (TRP1), orm2::TRP1 + pRS413 + ORM1/416 (pVB23) | This study | OSY1163 |
| SEY6210.1 MATa lcb1::TRP1 +pRS413 +pOS277 (3x-FLAG-LCB1-Δ50-85/416) | This study | OSY932 |
| **Oligonucleotides** |  |  |
| See Tab. S2 for used oligonucleotides |  |  |
| **Recombinant DNA** |  |  |
| pRS406-GAL1-ORM1-3A::URA3 | Schäfer et al., 2023<sup>21</sup> | FF457 |
| pRS404-GAL1-TSC3::TRP1 | Schäfer et al., 2023<sup>21</sup> | FF456 |
| pRS403-GAL1-SAC1::HIS3 | Schäfer et al., 2023<sup>21</sup> | FF476 |
| pRS405-GAL1-LCB2::LEU2 | Schäfer et al., 2023<sup>21</sup> | FF475 |
| pRS406-GAL1-3xFLAG-LCB1::URA3 | Schäfer et al., 2023<sup>21</sup> | FF474 |
| pRS405-GAL1-ORM2-3A::LEU2 | This study | FF478 |
| pRS405-GAL1-ORM2-3D::LEU2 | This study | FF622 |
| pRS406::URA3 | Sikorski and Hieter, 1989<sup>49</sup> | N/A |
| pRS406-ALFA-Lcb1::URA3 | Schäfer et al., 2023<sup>21</sup> | FF391 |
| pRS406-ALFA-*lcb1ΔTM1* | This study | FF535 |
| pRS413::HIS3 | Sikorski and Hieter, 1989<sup>49</sup> | N/A |
| pRS414::TRP1 | Sikorski and Hieter, 1989<sup>49</sup> | N/A |
| pRS415::LEU2 | Sikorski and Hieter, 1989<sup>49</sup> | N/A |
| pRS416::URA3 | Sikorski and Hieter, 1989<sup>49</sup> | N/A |
| pRS415-ALFA-LCB1::LEU2 | This study | pVN003 |
| pRS416-ORM1::URA3 | This study | pVB023 |
| pRS416-ORM1-3A::URA3 | This study | pOS274 |
| pRS416-ORM1-3D::URA3 | This study | pOS275 |
| pRS416-ORM2::URA3 | Schmidt et al., 2019<sup>35</sup> | pVB017 |
| pRS416-ORM2-3A::URA3 | This study | pOS321 |
| pRS416-ORM2-3D::URA3 | This study | pOS322 |
| pRS416-3xHA-ORM1::URA3 | Schmidt et al., 2019<sup>35</sup> | pOS127 |
| pRS416-3xFLAG-*LCB1*Δ50-85::URA3 | This study | pOS277 |
| **Software and algorithms** |  |  |
| Affinity Designer v.1.10.6 | Serif (Europe) Ltd | https://affinity.serif.com/de/designer/ |
| Perseus v.2.0.10 and v.2.0.11 | Tyanova et al., 2016<sup>50</sup> | https://maxquant.net/perseus/ |
| MaxQuant v2.4.2.0; v2.4.4.0; v.2.4.9.0 | Cox and Mann, 2008<sup>51</sup> | https://www.maxquant.org/ |
| R v.4.3.1 | The R Foundation | https://www.r-project.org/ |
| ImageJ v.1.53e | National Institutes of Health | https://ImageJ.nih.gov/ij/ |
| LigPlot<sup>+</sup> (v.2.2) | Laskowski and Swindells, 2011<sup>52</sup> | https://www.ebi.ac.uk/thornton-srv/software/LigPlus/ |
| AlphaFold (v.2.3.0) | Jumper et al., 2021<sup>53</sup> | https://github.com/google-deepmind/alphafold |
| UCSF ChimeraX (v.1.15) | Pettersen et al., 2004<sup>54</sup> | https://www.cgl.ucsf.edu/chimera/ |
| ChimeraX (v.1.5) | Meng et al., 2023<sup>55</sup> | https://www.cgl.ucsf.edu/chimerax/ |
| Phenix (v.1.19) | Liebschner et al., 2019<sup>56</sup> | https://phenix-online.org/ |
| ISOLDE (v.1.6) | Croll, 2018 | https://tristanic.github.io/isolde/ |
| Coot (v.0.9) | Emsley and Cowtan, 2004<sup>57</sup> | https://www2.mrc-lmb.cam.ac.uk/personal/pemsley/coot/ |
| MOLE (v.2.5) | Sehnal et al., 2013<sup>58</sup> | https://mole.upol.cz/ |
| PDBePISA | Krissinel and Henrick, 2007<sup>59</sup> | https://www.ebi.ac.uk/pdbe/pisa/ |
| EPU (v.2.9) | Thermo Fisher |  |
| cryoSPARC (v4) | Punjani et al., 2017<sup>60</sup> | https://cryosparc.com/ |
| PaSER (v.2023b) | Bruker |  |
| Affinity Photo v.1.10.1.1142 | Serif | RRID:SCR_016951 |
| Affinity Designer v. 1.10.1.1142 | Serif | RRID:SCR_016952 |
| Edge | Vilber Lourmat | Version 18.02 |
| **Other** |  |  |
| C-flat Holey Carbon (CF-1.2/1.3-3Cu-50) | Protochips | TEM-CF313-50 |

### Resource availability

#### Lead contact

Further information and requests for resources and reagents should be directed to and will be fulfilled by the lead contact, Florian Fröhlich.

#### Materials availability

Yeast strains and plasmids generated in this study are available upon request.

#### Data and code availability

- All density maps and models have been deposited in the Electron Microscopy Data Bank and the PDB. The PDB IDs are 8QOF for ScSPTOrm2-Dimer and 8QOG for ScSPTOrm2-Monomer. The respective EMDB IDs are EMD-18536, EMD-18537, and EMD-18533 for ScSPTOrm2-Dimer-masked.
- Mass spectrometric raw data have been deposited to the ProteomeXchange Consortium (http://proteomecentral.proteomexchange.org) via the PRIDE repository: PXD048754
- This paper does not report original code.
- Any additional information required to reanalyze the data reported in this work paper is available from the lead contact upon request.

#### Experimental model and study participant details

All yeast strains used in this study were generated from *S. cerevisiae* SEY6210.1, SEY6210 and W303 genetic background. A complete list of yeast strains and genotype information is provided in the key resources table. The list of oligos used in this study is provided in Tab. S2

#### Method details

##### Yeast strains

For yeast strains, plasmids and oligonucleotides used in this study, see Key resources table and Tab. S2. For purifications, subunits of the SPOTS complex were expressed under the control of the *GAL1* promoter using integrative plasmids and Lcb1 was internally 3xFLAG-tagged after P9, as previously reported^19^.

##### Purification of 3xFLAG tagged SPOT-Orm2 complex from *S. cerevisiae*

Purification of the 3xFLAG tagged SPOTS complex from yeast cells was performed as previously described^21^. Yeast cells, grown in yeast peptone (YP) medium containing 2% galactose (v/v), were washed in lysis buffer (50 mM HEPES-KOH (pH 6. 8), 150 mM KOAc, 2 mM MgOAc, 1 mM CaCl_2_, 200 mM sorbitol) and resuspended in lysis buffer containing 1 mM phenylmethylsulfonyl fluoride (PMSF) and 1x FY protease inhibitor mix (Serva) in a 1:1 ratio (w/v). Cells frozen dropwise in liquid nitrogen were pulverized in 15 × 2-minute cycles at 12 CPS in a 6875D Freezer/Mill Dual-Chamber Cryogenic Grinder (SPEX SamplePrep) and thawed in lysis buffer containing 1 mM PMSF and 1x FY. The lysate was cleared by two centrifugation steps at 1,000 *g* at 4 °C for 20 min and microsomal membranes were pelleted at 44,000 *g* at 4 °C for 30 min. Cells were resuspended in lysis buffer followed by the addition of IP buffer (50 mM HEPES-KOH, pH 6.8, 150 mM KOAc, 2 mM MgOAc, 1 mM CaCl_2_, 15% glycerol) containing 1% glyco-diosgenin (GDN) supplemented with protease inhibitors. Membrane proteins were solubilized for 1.5 hours at 4 °C nutating and non-solubilized material was removed by centrifugation for 30 min at 44,000 *g* at 4 °C. Cleared lysate was added to α-FLAG resin (Sigma Aldrich) and nutated for 45 min at 4 °C. Beads were washed twice with 20 ml IP buffer containing 0.1% and 0.01% GDN, respectively. SPOTS complex was eluted twice in IP buffer containing 0.01% GDN for 45 min and 5 min at 4 °C with 3xFLAG peptide. After concentrating in a 100 kDa Amicon Ultra centrifugal filter (Merck Millipore), size exclusion chromatography (SEC) was performed with the concentrated eluate on a Superose 6 Increase 5/150 column (Cytiva) using the ÄKTA go purification system (Cytiva). Peak fractions were collected and concentrated followed by cryo-EM and mass spectrometric analysis.

##### Cryo-EM sample preparation and data acquisition

Sample quality was inspected by negative-stain electron microscopy as previously described (Fig. S 1D) ^61^. Micrographs of the negatively-stained sample were recorded manually on a JEM2100plus transmission electron microscope (Jeol), operating at 200 keV and equipped with a Xarosa CMOS (Emsis) camera at a nominal magnification of 30,000, corresponding to a nominal pixel size of 3.12 Å per pixel. For cryo-EM, the sample was concentrated to 10 mg/ml. C-flat grids (Protochips; CF-1.2/1.3-3Cu-50) were glow-discharged, using a PELCO easiGlow device at 15 mA for 45 s and 3 µl of the concentrated sample were immediately applied and plunge frozen in liquid ethane, using a Vitrobot Mark IV (Thermo Fisher) at 100 % relative humidity, 4 °C. The dataset was collected using a Glacios microscope (Thermo Fisher), operating at 200 keV and equipped with a Selectris energy filter (Thermo Fisher) with a slit of 10 eV. Movies were recorded with a Falcon 4 direct electron detector (Thermo Fisher) at a nominal magnification of 130,000 corresponding to a calibrated pixel size of 0.924 Å per pixel, and the data was saved in the electron-event representation (EER) format. The dose rate was set to 5.72 e-per pixel per second and a total dose of 50 e-per Å2. 15,822 movies were collected automatically, using EPU software (v.2.9, Thermo Fisher) with a defocus range of −0.8 to −2.0 µm.

##### Cryo-EM image processing

The SPOT dataset was processed in cryoSPARC (v.4), and the processing workflow is presented in Fig. S 2. Movies were preprocessed with patch-based motion correction, patch-based CTF estimation and filtered by the CTF fit estimates using a cut-off at 5 Å in cryoSPARC live (v.4), resulting in a remaining stack of 13,168 micrographs (representative micrograph given in Fig. S 2). Well-defined 2D classes were selected and used for subsequent rounds of template-based 2D classification. 1,208,193 particles were extracted in a box of 432 pixels, and Fourier cropped to 216 pixels. Previously generated ab initio 3D reconstructions from live processing were used for several rounds of heterogeneous refinement, which resulted in two distinct, well-defined reconstructions. Both reconstructions were processed separately. Processing of the SPOT-dimer-complex: Classes corresponding to the newly termed SPOT-dimer-complex were subjected to non-uniform refinement, and 3D-aligned particles were re-extracted without binning. Additional rounds of heterogeneous refinement following non-uniform refinement and local refinement with C2 symmetry applied resulted in a stack of 159K particles and a consensus map with 3.3 Å resolution. To further improve the map quality, particles were symmetry expanded by using a C2 point group, thereby aligning signals coming from both protomers onto the same reference map. Subsequent signal subtraction and local refinement results were subjected to 3D classification in PCA mode focused around the protomer. Additional local CTF correction and local refinement of 152K symmetry-expanded particles yielded a focused map with an overall resolution of 2.8 Å. Processing of the SPOT-monomer: An additional well-resolved 3D class was identified and processed separately. 328K particles were cleaned through 2D classification to remove the remaining non-protomer classes. A stack of 170K particles was subjected to heterogeneous refinement, and the remaining 142K particles were further refined through 3D-classification in PCA-mode, non-uniform and local refinement. This resulted in a final map with a global resolution of 3.1 Å. Reported B-factors resulted from un-supervised auto-sharpening during refinement in cryoSPARC. To aid model building, unsharpened half-maps were subjected to density modification within Phenix phenix.resolve_cryo_em^56^.

##### Model building and refinement

Initial atomic models were generated using the AlphaFold2 prediction of the individual subunits, which were placed in the individual maps and fitted as rigid bodies through UCSF Chimera^54^. The structure was manually inspected in Coot (v.0.9)^57^ and iteratively refined using phenix.real_space_refine within Phenix (v.1.19). Calculation of compositional heterogeneity of the reconstructions was performed with OccuPy^38^. Validation reports were automatically generated by MolProbity^62^ within Phenix. All density maps and models have been deposited in the Electron Microscopy Data Bank and the PDB. The PDB IDs are 8QOF for *Sc*SPTOrm2-Dimer and 8QOG for *Sc*SPTOrm2-Monomer. The respective EMDB IDs are EMD-18536, EMD-18537, and EMD-18533 for ScSPTOrm2-Dimer-masked. Structures were visualized with ChimeraX^54^ and protein interactions were analyzed with the help of PDBe PISA^59^. 2D ligand-protein interaction diagrams in Fig. S 5 were calculated in LigPlot+^52^. An overview of the local density fit is given in Fig. S 4.

##### Computational prediction of protein structures

The model for full-length SPOT-Orm2 (ScLcb1: P25045, ScLcb2: ScP40970, Orm2: ScQ06144 and Tsc3: Q3E790) was predicted using the multimer-implementation of AlphaFold2^53^. The template date was set to 2020-05-14 and predictions were run on a local workstation. Validation parameters of the prediction, including the per-residue model confidence score pLDDT and the PAE (prediction aligned error matrix) were visualized using a custom python script. Phosphorylations were modeled with PyTMs^63^.

##### Pulldown experiments

Independent biological triplicates (n=3) were inoculated from an overnight pre-culture in 200 ml YPD and grown to exponential growth phase at 30 °C. For each sample, an equal amount of cells was harvested at 2,272 *g* at 4 °C for 5 min and snap frozen in liquid nitrogen. Cell lysis was performed with 500 μl glass beads in 500 μl pulldown buffer (20 mM HEPES pH 7.4, 150 mM KOAc, 5% glycerol, 1% GDN, Roche Complete Protease Inhibitor Cocktail EDTA free, Roche) using a FastPrep (MP biomedicals). The cleared lysate (1,000 *g*, 10 min, 4 °C) was incubated for 30 min at 4 °C on a turning wheel to solubilize membrane proteins. Unsolubilized material was removed at 21,000 *g* at 4 °C for 10 min and supernatants were incubated with 12.5 μl of pre-equilibrated ALFA beads (NanoTag Biotechnologies) on a turning wheel at 4 °C. Beads were washed twice with pulldown buffer followed by four washing steps with wash buffer (20 mM HEPES pH 7.4, 150 mM KOAc, 5% glycerol). Bound proteins were digested and prepared according to the iST Sample Preparation Kit (PreOmics). Dried peptides were resuspended in 10 μl LC-Load, and 2 μl were used to perform reversed-phase chromatography on a Thermo UltiMate 3000 RSLCnano system connected to a TimsTOF HT mass spectrometer (Bruker Corporation, Bremen) through a Captive Spray Ion source. Separation of peptides was performed on an Aurora Gen3 C18 column (25 cm x 75um x 1.6um) with CSI emitter (Ionoptics, Australia) at 40 °C followed by their elution via a linear gradient of acetonitrile from 10–35% in 0.1% formic acid for 44 min at a constant flow rate of 300 nl/min following a 7 min increase to 50%, and finally, 4 min to reach 85% buffer B. Eluted peptides were directly electro sprayed into the mass spectrometer at an electrospray voltage of 1.5 kV and 3 l/min Dry Gas. The MS settings of the TimsTOF were adjusted to positive ion polarity with a MS range from 100 to 1700 m/z Resulting data were analyzed using MaxQuant (V2.4.9.0, www.maxquant.org)^51,64^ and Perseus (V2.0.7.0, www.maxquant.org/perseus)^50^. Significance lines in the volcano plot of the Perseus software package corresponding to a given FDR were determined by a permutation-based method^65^. The mass spectrometry proteomics data have been deposited to the ProteomeXchange Consortium via the PRIDE partner repository^66^.

##### LCB and ceramide analysis of purified SPOTS complex and whole cell lysate

LCBs and ceramide were extracted and measured from 10 µg purified proteins or from an equivalent of 150 μg protein from whole cell lysate in biological independent quadruplicates (n = 4). For whole cell lysate extraction, cells were grown in YPD to exponential growth phase and 150 mM ammonium formate was added to cell lysate. Ceramide (Sphingosine d17:1, CER d17:1/24:0; Avanti) was added as an internal standard and lipids were extracted with 2:1 chloroform/methanol as described previously^67,68^. Dried lipid films were dissolved in a 65:35 (v/v) mixture of Buffer A (50:50 water/acetonitrile, 10mM ammonium formate, and 0.1 % formic acid) and B (88:10:2 2-propanol/acetonitrile/water, 2mM ammonium formate and 0.02 % formic acid). Dihydrosphingosine 18:0 (DHS; Avanti Polar Lipids), phytosphingosine 18:0 (PHS; Avanti Polar Lipids) and phytoceramide t18:0/24:0 (Avanti Polar Lipids 567 Lipids/Cayman) were used for an external standard curve. Samples were analyzed on an Accucore C30 LC column (150mm× 2.1mm 2.6 μm Solid Core; Thermo Fisher Scientific) connected to a Shimadzu Nexera HPLC system and a QTRAP 5500 LC-MS/MS (SCIEX) mass spectrometer. For the gradient, 40 % of Buffer B was used for 0.1 min, then increased to 50 % over 1.4 min followed by an increase of Buffer B to 100 % over 1.5 min. After 1 min of 100 % Buffer B, B was decreased to 40 % for 0.1 min and kept at 40 % until the end of the gradient. A constant flow rate of 0.4 ml/min was set and a total analysis time of 6 min was used. As an injection volume, 1 or 2 µl were used. The MS data were measured in positive ion, scheduled MRM mode without detection windows (Tab. S3). The SciexOS software was used for peak integration. The concentrations of all lipid species, expressed in pmol/µg protein, were determined using the external standard curve.

##### Colorimetric-based enzymatic assay

SPT activity was determined by monitoring released CoA-SH from the SPT-catalyzed condensation of palmitoyl-CoA and L-serine coupled to an enzymatic assay using the Coenzyme A Assay Kit from Sigma-Aldrich. All assays were performed on a 200 µl scale in IP buffer with 0.008 % GDN, 15 mM L-serine, 100 µM palmitoyl-CoA, and 30 µM PLP. The reaction was initiated by the addition of 1 µg purified protein. As a control, the SPT-specific inhibitor myriocin was included at a final concentration of 100 µM and methanol (MeOH) was added to all other samples since myriocin was dissolved in MeOH. The reactions were incubated for 1 h at RT followed by their deproteinization with a 3 kDa MWCO concentrator (Merck Millipore). CoA levels were measured in a 96-well plate according to the Coenzyme A Assay Kit protocol (Sigma-Aldrich, MAK034). CoA concentration was determined by measuring the final colorimetric product at 570 nm in a SpectraMax iD3 Multi-Mode microplate reader. Absorbances of the protein-free samples were subtracted from the absorbances of the protein-containing samples to obtain the final corrected absorbances.

##### Spotting assays

Cells were inoculated in YPD from an overnight pre-culture and grown to exponential growth phase. Serial dilutions were prepared and spotted onto YPD plates and YPD plates containing the SPT-specific inhibitor myriocin. Plates were incubated at 30 °C for 3 days.

##### Preparation of whole cell protein extracts for Western blot

To prepare whole cell lysates, proteins were extracted by a modified alkaline extraction protocol (Schmidt et al., 2019). Cells (4 OD_600nm_) were harvested by centrifugation (5 min, 4000 rpm, 4°C) and washed in ice cold water with phosphatase inhibitors (10mM NaF, 10 mM beta-glycerol phosphate, PhosStop (Roche, 1 tablet per 100 mL). After centrifugation (3 min, 13000 rpm, 4°C) cells were re-suspended in 0.1M NaOH with the same phosphatase inhibitors and incubated at room temperature for 5 min. After centrifugation (3 min, 13000 rpm, 4°C) pellets were re-suspended in Lämmli sample buffer, denatured (95°C, 12 min), and cell debris was removed by centrifugation (3 min, 13000 rpm, 4°C).

##### Western Blot analysis and immunodetection

Protein extracts dissolved in Lämmli sample buffer were separated by SDS-PAGE (Biorad Mini Protean) and transferred to PVDF membranes by semi-dry or wet electro-blotting. Antibodies used in this study are listed in the key resources table.

##### Phos-tag SDS PAGE

Phos-tag gels (50 µM Phos-tag acrylamide (Wako); 100 µm MnCl_2_) were prepared according to the manufacturer’s specifications. Gels were run in a standard Lämmli electrophoresis buffer at 200V, 40 mA for 1.5 hours, afterwards rinsed in Western blot transfer buffer with 10 mM EDTA for 20 minutes and equilibrated in transfer buffer, followed by standard wet electroblotting to PVDF membranes.

##### Native immunoprecipitation of ALFA-Lcb1 or FLAG-Orm1

Logarithmically growing cells (1000 OD_600nm_) were harvested by centrifugation, and pellets were frozen in liquid nitrogen. Frozen cells were ground in a 6770 freezer mill (SPEX Sample Prep) with liquid nitrogen cooling. 200 mg yeast powder was resuspended in 1 ml lysis buffer (50 mM HEPES/NaOH, pH 6.8; 150 mM NaOAc, 2mM Mg(OAc)_2_; 1mM CaCl_2_; 15% glycerol) supplemented with 2% GDN (Avanti polar lipids); protease and phosphatase inhibitor (10 µg/ml Leupeptin; 1 µg/ml Aprotinin; 1 µg/ml Pepstatin A; 0.4 mM PEFA-Bloc; 1x yeast protease inhibitor mix (Sigma Aldrich); 1x Complete EDTA free (Roche); 1x PhosStop (Roche)) and incubated rotating at 4°C for 1h. The lysate was mixed with an equal volume (1 ml) of lysis buffer with 0.1% GDN and incubated rotating at 4°C for 30 min. Unsolubilized material was removed by centrifugation (15.000 g; 20 min; 4°C). The supernatant (1.6 ml) was added to 15 µl equilibrated anti-FLAG magnetic beads (Sigma Aldrich Cat. # M8823; RRID:AB_2637089) or 10 µl magnetic anti-ALFA ST selector beads (NanoTag Cat.# N1516), and rotated for 3h at 4°C. Beads were recovered with a magnetic rack and washed 5 times for 10 minutes with 2 ml lysis buffer containing 0.1% GDN. FLAG beads were eluted twice with 150 µl lysis buffer supplemented with 0.2% GDN and 0.5 mg/ml 3xFLAG peptide (Sigma Aldrich) at 4°C for 15 minutes. Elutions were pooled, supplemented with 100 µl 4xSDS PAGE sample buffer and denatured (95°C, 10 min). ALFA beads were eluted with 150 µl lysis buffer containing 0.1% GDN mixed with 50 µl 4x SDS PAGE sample buffer by boiling at 95°C for 10 minutes.

##### Induced loss of *URA3* plasmids by growth on 5-FOA

Heterozygous diploid strains were generated by mating adequate precursor strains and transformed with pVB023 (*ORM1* in pRS416) or pOS277 (*3xFLAG-LCB1Δ50-85* in pRS416). Strains were subsequently sporulated, the ensuing tetrads dissected with a Singer MSM400 micromanipulator, and the desired haploid genotypes were obtained by auxotrophy and mating type analysis. Haploid cells containing the cover plasmids were grown in liquid culture (170 rpm; 26°C) into exponential phase in selective medium lacking uracil.

For spot assays, 1 OD cells was harvested by centrifugation, resuspended in 1 ml sterile water and diluted to 0.1 and 0.01 OD/ml. 10 µl drops were placed on synthetic complete medium (containing uracil) supplemented with 0.75 mg/ml 5-fluoroacetic acid (5-FOA) where indicated and incubated at 26°C for up to 7 days.

For Western blot and lipidomics analysis, exponentially growing cells were diluted to OD 0.1 into synthetic complete liquid medium supplemented with 0.75 mg/ml 5-FOA and grown at 26°C, 170 rpm. After 24 hours, cultures were diluted to OD 0.1 in 50 ml synthetic complete + 0.75 mg/ml 5-FOA. Cells were kept in exponential growth for another 24 hours before cells were harvested (4 OD for whole cell extracts, 25 OD for lipid extraction).

#### Quantification and statistical analysis

Data analysis of all the experimental results such as mean value and standard deviations was performed in Microsoft Excel. Data were analyzed in R Software (v.4.3.1) using one-way ANOVA with Tukey’s multiple-comparison test (\**P* < 0.05, \*\**P* < 0.01, \*\*\**P* < 0.001). Exact *P*-values are shown in Tab. S4.

## Acknowledgements

We thank Teresa Dunn and Robbie Loewith for antisera. This work was funded by the DFG (SFB1557 P6 to FF, P11 to AM, FR 3647/2-2 and FR 3647/4-1, INST 190/201-1, INST.190/196-1 FUGG and BMBF 01ED2010 to AM. JHS is supported by a fellowship from the Friedrich Ebert foundation. OS receives funding from the Austrian Science Fund (FWF P 36187-B). DT receives funding from the Austrian Science Fund (FWF P 32161, P 34907).

## Author contributions

C.K., J-H. S, O.S., A.M. and F.F. designed the project. C.K., J-H. S. and O.S. performed and analyzed most of the experiments. S.W. and B.M.S. performed and analyzed all lipidomics experiments. C.K., S. W. and F.F. performed and analyzed proteomics experiments. J-H.S., D.J., K.P. and A.M. generated and evaluated all cryo-EM data. C.K. performed all protein purifications. A.M., D.T., O.S. and F.F. supervised experiments and analyzed data. C.K. J-H.S., A.M., O.S. and F.F. prepared the manuscript. All authors discussed the results, contributed to the manuscript, and agreed on the content of the paper.

