## Supporting Information for "The structure of the Orm2-containing serine palmitoyltransferase complex reveals distinct inhibitory potentials of yeast Orm proteins"

### Correspondence e-mail:

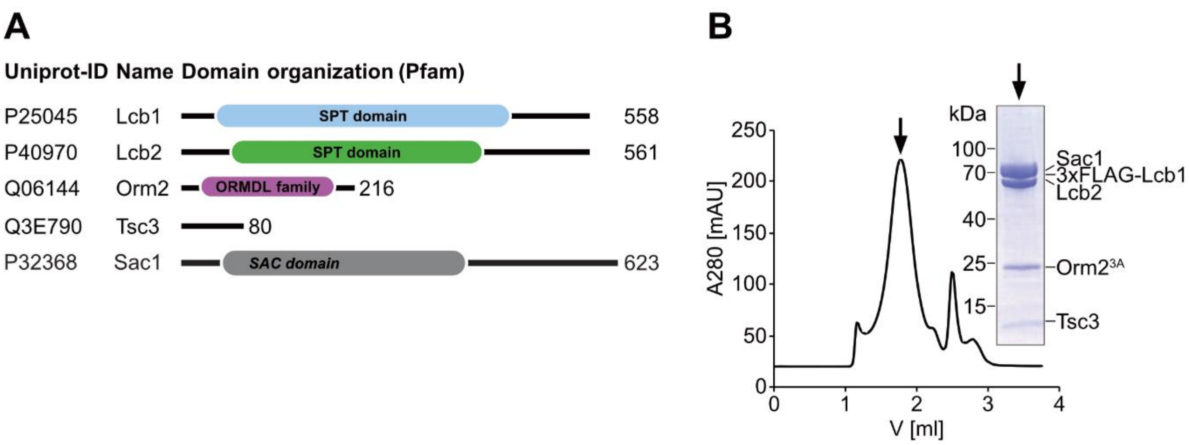


C

| Gene name | Unique peptides | Sequence coverage | Intensity |
|---|---|---|---|
| *LCB1* | 14 | 37.6 | $3.8 \cdot 10^{10}$ |
| *LCB2* | 17 | 34.8 | $3.8 \cdot 10^{10}$ |
| *TSC3* | 2 | 30 | $7.2 \cdot 10^{9}$ |
| *SAC1* | 33 | 63.2 | $6 \cdot 10^{10}$ |
| *ORM2* | 3 | 28.7 | $2.9 \cdot 10^{8}$ |

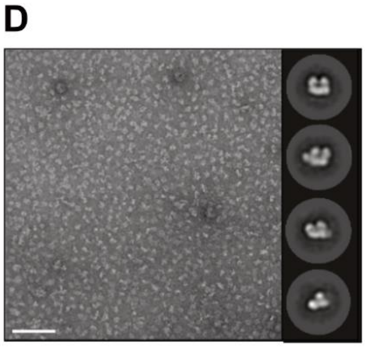


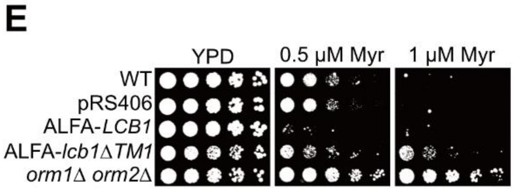


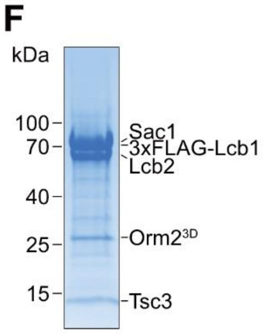


**Fig. S1 A,** Pfam-based domain annotation of SPOT-Orm2 subunits. **B**, Size-exclusion chromatography profile of GDN-solubilized SPOT-Orm2$^{3A}$ complex and Coomassie-blue stained SDS-PAGE gel of purified SPOT-Orm2$^{3A}$ complex from highlighted fraction after concentrating. **C**, Summary of the mass spectrometric analysis of SPOT-Orm2$^{3A}$ in B. **D**, Representative micrograph and 2D class averages from negative-stain TEM. 100 nm scale bar. **E**, Serial dilutions of control cells, *lcb1Δ* cells expressing ALFA-*LCB1* or ALFA-*lcb1ΔTM1* and *orm1Δ orm2Δ* cells on YPD plates (control) and YPD plates containing 0.5 µM and 0.75 µM of the SPT-specific inhibitor myriocin. **F**, Coomassie-blue stained SDS-PAGE gel of GDN-solubilized SPOT-Orm2$^{3D}$ complex.

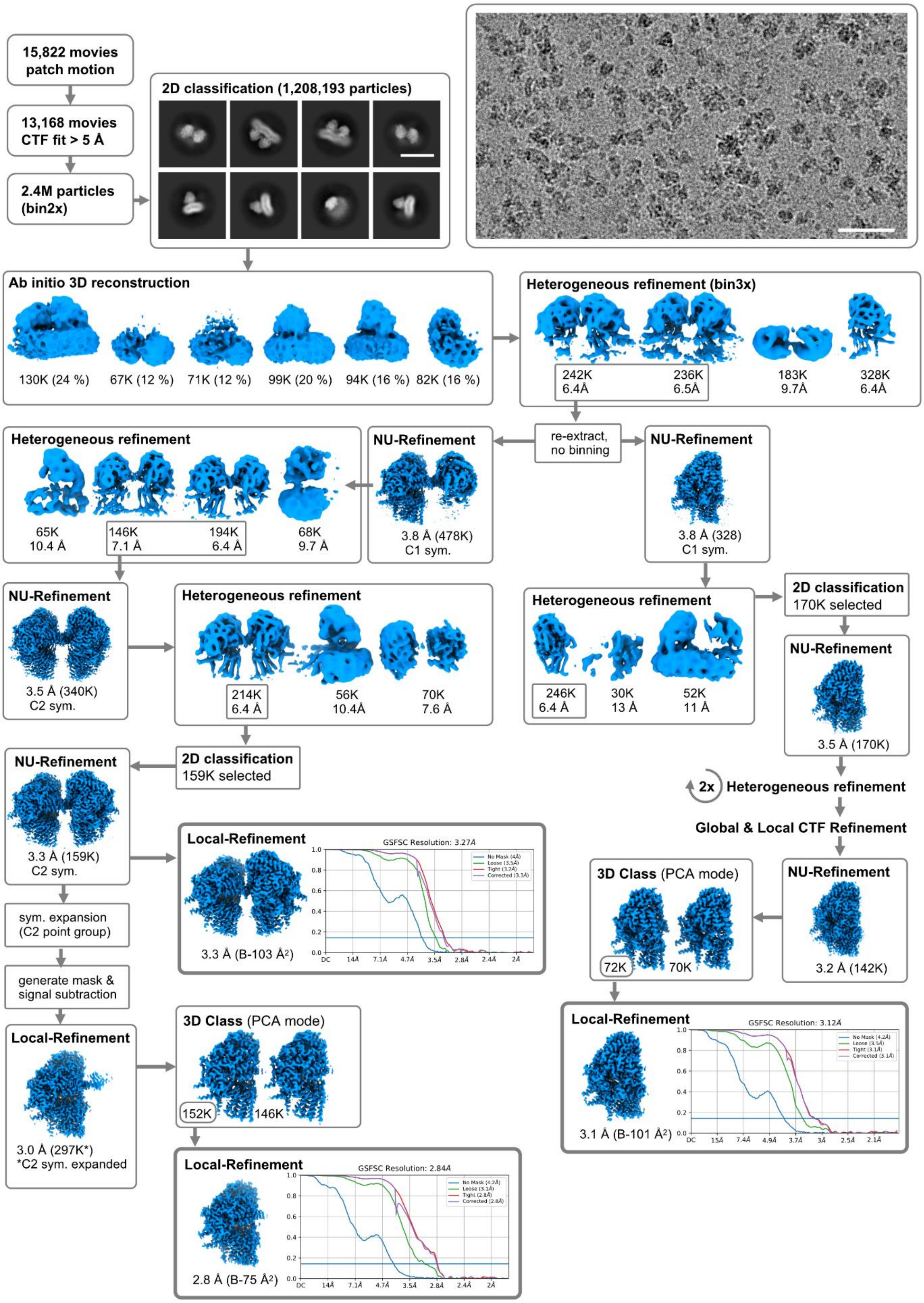


**Fig. S2** Cryo-EM analysis of SPT-Orm2 complexes: Processing workflow for SPOT-dimer and SPOT-monomer. All processing steps were performed in cryoSPARC. Representative cryo-EM micrograph and 2D-class averages. 100 nm scale bar in micrograph and 20 nm scale bar in 2D class-averages.

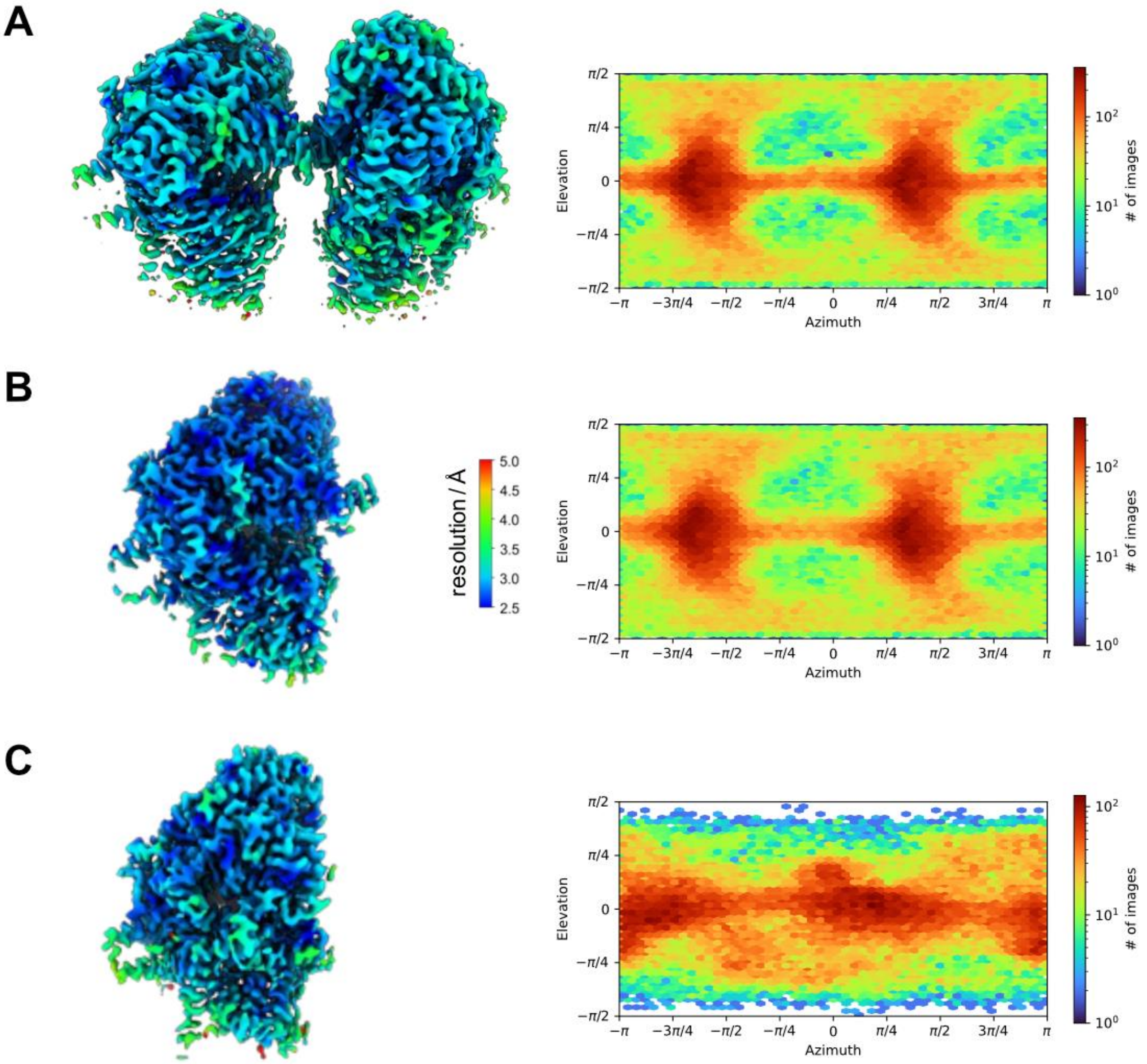


**Fig. S3** Cryo-EM map evaluation: Local resolution estimation and angular particle distribution of SPOT-dimer (level 0.18, **A**), masked SPOT-dimer (level 0.18, **B**), SPOT-monomer (level 0.18, **C**).

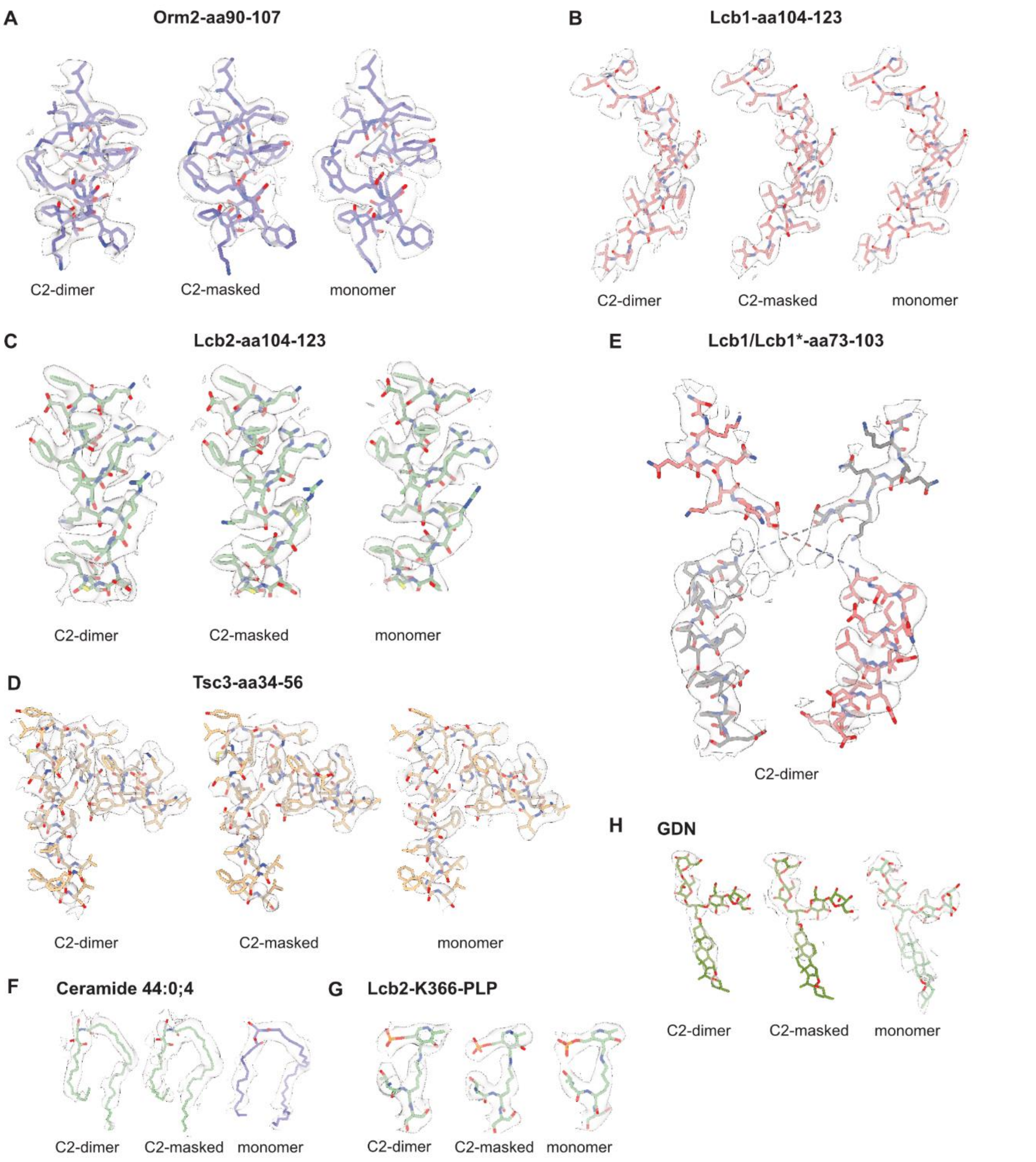


**Fig. S4** Local cryo-EM density map quality of SPOT complexes: Side-by-side comparison of selected residues and ligands within all cryo-EM density maps of **A** Orm2, **B** Lcb1, **C** Lcb2, **D** Tsc3, **E** crossover of Lcb1-TM1, **F** Ceramide 44:0;4, **G** Lcb2-K366-PLP (pyridoxal phosphate) and **H** GDN (glyco-diosgenin). Contouring levels set to 0.16 for all densities.

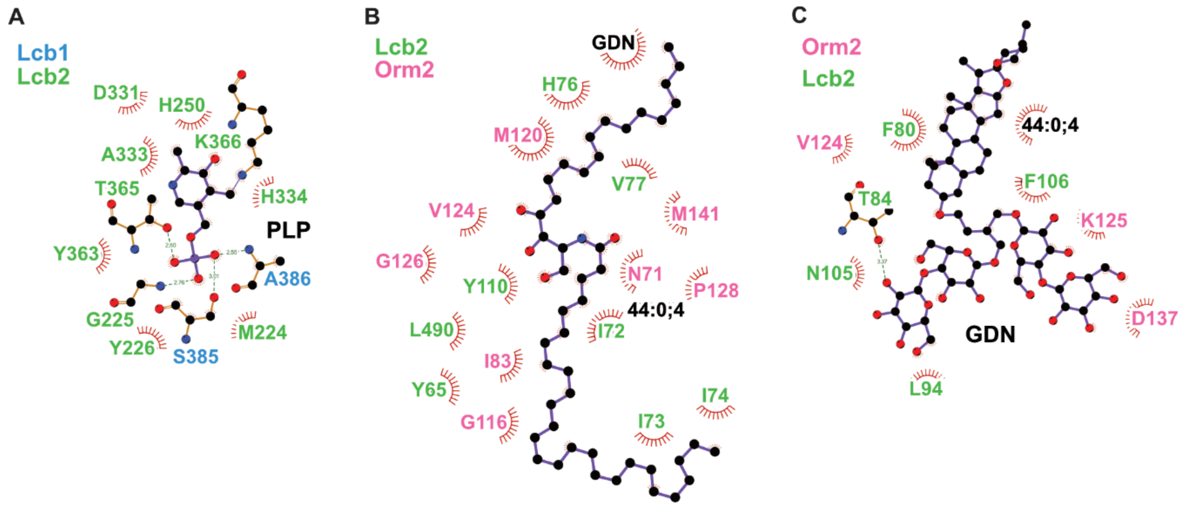


**Fig. S5** Ligand interaction diagram of SPOT complexes. **A**, 2D ligand interaction diagram of internal aldimine between PLP and Lcb2-K366, **B**, ceramide 44:0;4, and **C**, glyco-diosgenin (GDN). Key interacting residues are shown as ball and sticks with polar contacts given as green dotted lines. Diagrams were calculated and visualized with LigPlot+[1].

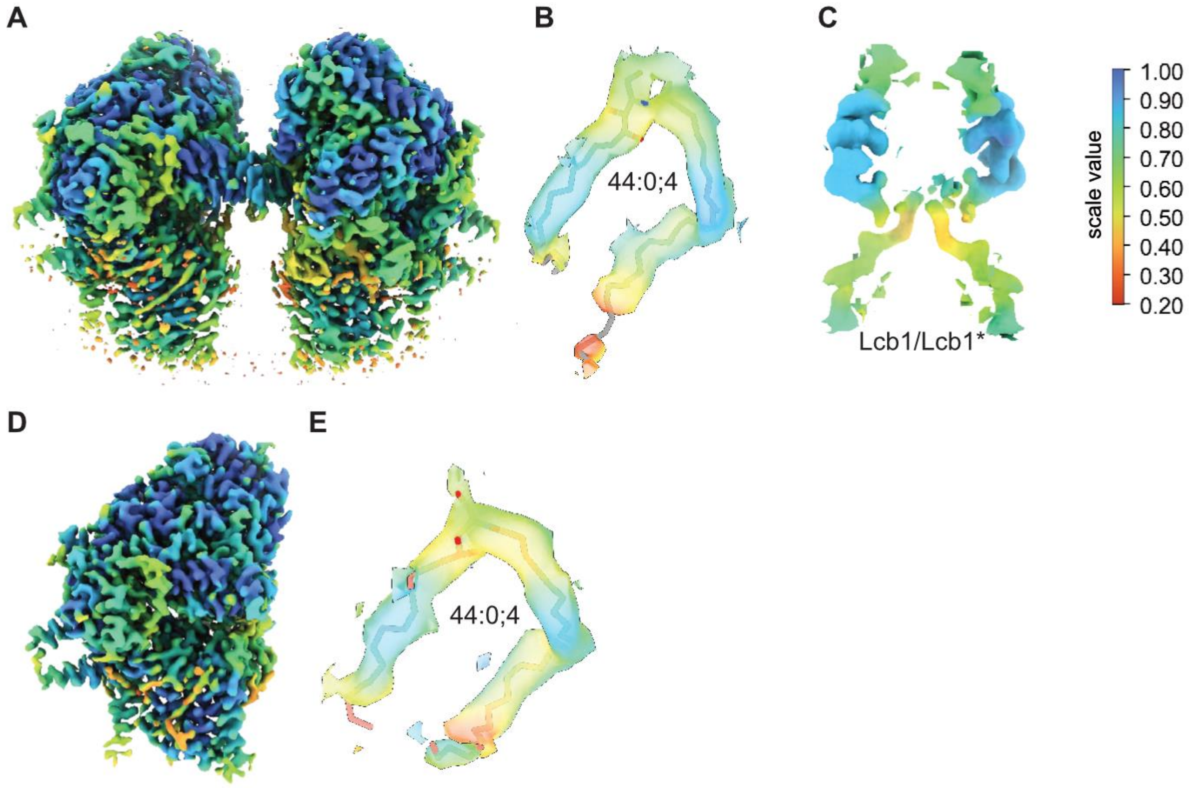


**Fig. S6** Estimation of compositional heterogeneity. Scale-value color-coded cryo-EM densities are shown. **A**, SPOT-dimer overview. **B**, SPOT-dimer ceramide 44:0;4. **C**, SPOT-dimer Lcb1/Lcb1* crossover region. **D**, SPOT-monomer overview. **E**, SPOT-monomer ceramide 44:0;4. All maps are contoured automatically by OccuPy[2]. Low compositional heterogeneity is shown in blue, high values are shown in red.

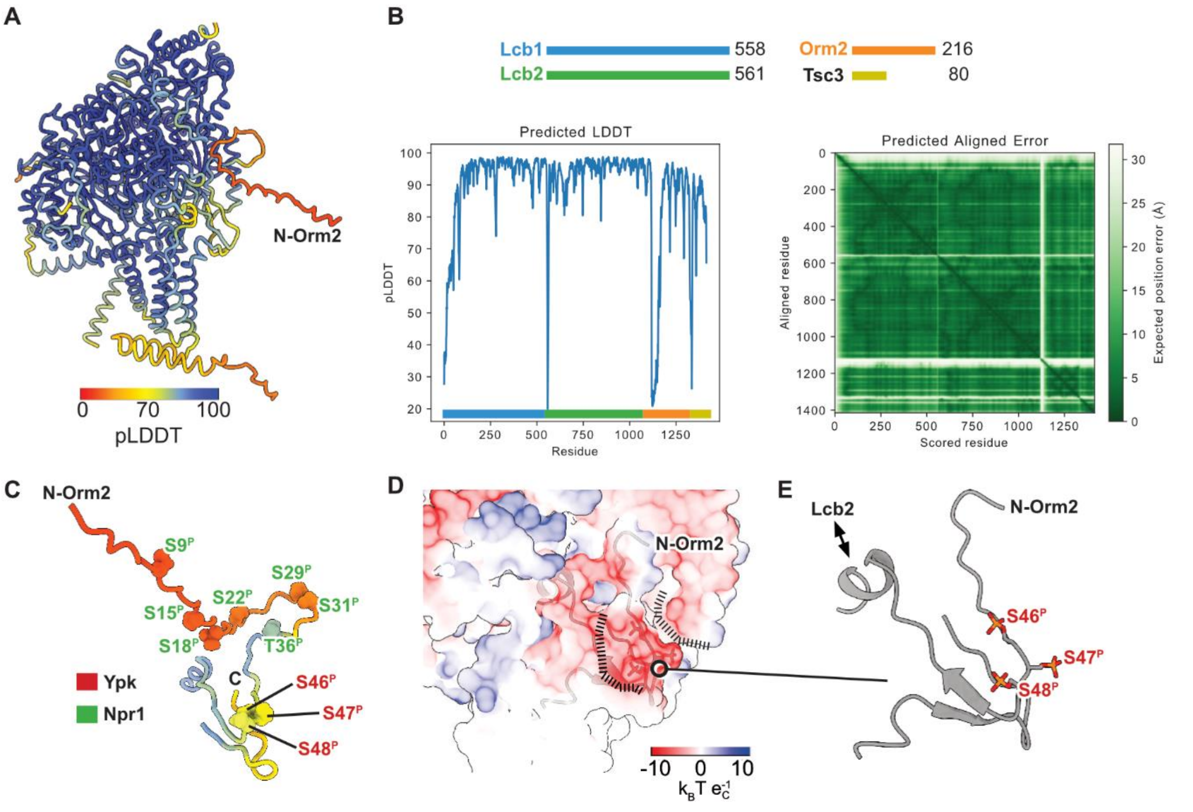


**Fig. S7 A**, AlphaFold2 prediction in multimer mode of SPOT-Orm2 monomer, color-coded by the predicted local distance difference test (pLDDT). The low-confidence N-terminus of Orm2 is annotated accordingly. **B**, Validation parameters of the prediction, including the pLDDT and the PAE (prediction aligned error matrix). The pLDDT plot is color-coded according to the different subunits. The prediction was run locally using AlphaFold v.2.1.02 in the multimer-mode (template date set to 2020-05-14). **C**, Phosphorylation sites of the N-terminal region of Orm2. Target sites for Ypk kinases in red and Npr1 kinase in green. Protein-backbone trace color-coded by pLDDT. **D**, Electrostatic surface potential focusing on the N-terminal region of Orm2. **E**, Cartoon-representation of N-terminal region of Orm2 with modelled phospho-serine.

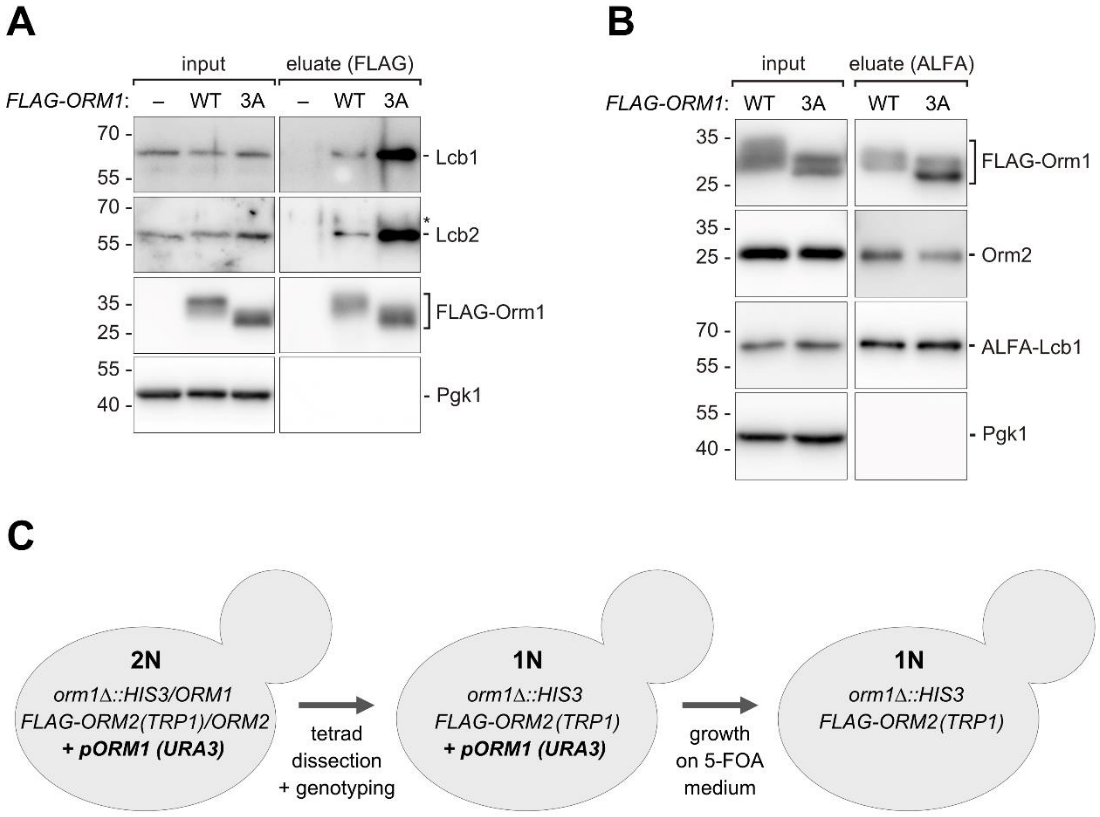


**Fig. S8 A**, Input and eluate fractions of a native FLAG immunoprecipitation from WT cells and cells chromosomally expressing *FLAG-ORM1* WT or the 3A mutant from the endogenous locus analyzed by SDS PAGE and Western blotting with the indicated antibodies. **B**, Input and eluate fraction of a native immunoprecipitation of ALFA-Lcb1 from cells chromosomally expressing *FLAG-ORM1* WT or the 3A mutant from the endogenous locus analyzed by SDS PAGE and Western blotting with the indicated antibodies. **C**, Schematic illustrating the procedure for the generation of cells expressing only a single SPOTS complex variant (containing either FLAG-Orm1 WT, FLAG-Orm1$^{3A}$, FLAG-Orm2 WT or FLAG-ORM2$^{3A}$) through induced loss of a plasmid encoding *ORM1* WT on 5-FOA medium. The cartoon describes the generation of the *orm1*Δ *FLAG-ORM2* WT strain. The other analyzed strains were generated accordingly.

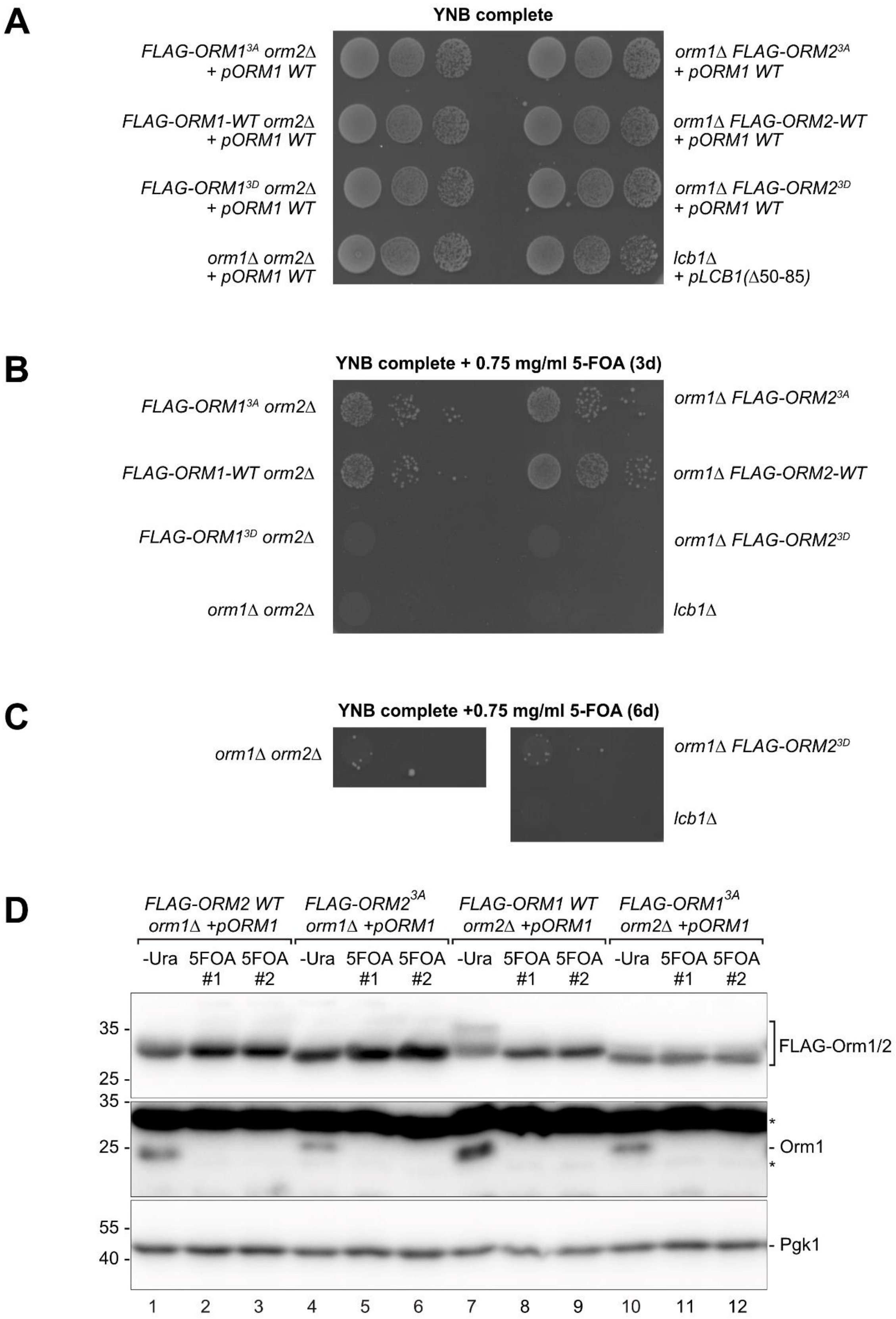


**Fig. S9 A**, Growth of the indicated strains (containing *ORM1* or *LCB1Δ50-85* cover plasmids with *URA3* selection markers) on synthetic complete medium. Plates were incubated for 3 days at 26°C. **B**, The same strains as in A) grown for 3 days at 26°C on synthetic complete plates supplemented with 0.75 mg/ml 5-FOA to induce loss of the cover plasmids. **C**, The indicated strains after 6 days at 26°C on synthetic complete plates supplemented with 0.75 mg/ml 5-

FOA. **D**, Whole cell protein extracts of the indicated strains before (-Ura) and after growth in synthetic complete liquid medium supplemented with 0.75 mg/ml 5-FOA for 48 hours (two independent 5-FOA cultures #1 and #2) were analyzed by SDS PAGE and Western blotting with the indicated antibodies. * indicates cross-reactions of the Orm1 antiserum.

**Tab. S1** Cryo-EM data collection, refinement and validation statistics.

*Saccharomyces cerevisiae* serine palmitoyltransferase (SPT) with Orm2, Lcb1, Lcb2 and Tsc3.

| | *Sc*SPT-Orm2 dimer (PDB 8QOF, EMD-18536) | *Sc*SPT-Orm2 dimer masked (EMD-18533) | *Sc*SPT-Orm2 monomer (PDB 8QOG, EMD-18537) |
|---|---|---|---|
| Data collection and processing | | | |
| Microscope | TFS Glacios | TFS Glacios | TFS Glacios |
| Voltage (keV) | 200 | 200 | 200 |
| Camera | TFS Falcon 4 | TFS Falcon 4 | TFS Falcon 4 |
| Energy filter | TFS Selectris | TFS Selectris | TFS Selectris |
| Slit width eV | 10 | 10 | 10 |
| Magnification (nominal) | 130,000 | 130,000 | 130,000 |
| Pixel size (Å/px) | 0.945 | 0.945 | 0.945 |
| Defocus range (µm) | 0.8-2.0 | 0.8-2.0 | 0.8-2.0 |
| Total exposure ($e^-$/Å$^2$) | 50 | 50 | 50 |
| Exposure rate ($e^-$/px/s) | 5.72 | 5.72 | 5.72 |
| Automation software | *EPU 2.9* | *EPU 2.9* | *EPU 2.9* |
| Processing software | *CryoSPARC* 4 | *CryoSPARC* 4 | *CryoSPARC* 4 |
| Micrographs collected* | 15,822 | 15,822 | 15,822 |
| Micrographs used* | 13,168 | 13,168 | 13,168 |
| Final particle images | 148,610* | 148,610* | 71,624 |
| Point-group symmetry parameters | *C2* | *C2* | *C1* |
| Resolution (global) (Å) FSC 0.134 | 3.3 | 2.8 | 3.1 |
| Map-sharpening *B* factor (Å$^2$) | – 103 | – 75 | – 101 |
| Map-sharpening method | Global *B* factor | Global *B* factor | Global *B* factor |
| Refinement package | *phenix.real_space_refine* | | phenix.real_space_refine |
| Model Composition | | | |
| Non-H atoms | 20,702 | | 10,444 |
| Protein Residues | 2,569 | | 1,297 |
| Model refinement | | | |
| Model-map scores | | | |
| CC (mask) | 0.79 | | 0.83 |
| CC (volume) | 0.80 | | 0.80 |
| FSC model (0.5) | 3.4 | | 3.5 |
| Av. grouped B factors (Å$^2$) | | | |
| Protein residues | 52.64 | | 61.21 |
| Ligands | 22.57 | | 24.18 |
| R.m.s.d. from ideal values | | | |
| Bond lenghths (Å) | 0.005 | | 0.004 |
| Bond angles (°) | 1.031 | | 0.672 |
| Validation | | | |
| *MolProbity* score | 1.42 | | 1.38 |
| CaBLAM outliers (%) | 2.46 | | 2.65 |
| Clashscore | 3.94 | | 3.23 |
| Poor rotamers (%) | 0.13 | | 0.35 |
| $C^\beta$ outliers (%) | 0.00 | | 0.16 |
| Ramachandran plot | | | |
| Favored (%) | 96.39 | | 96.12 |
| Allowed (%) | 3.61 | | 3.88 |
| Outliers (%) | 0.00 | | 0.00 |

* Single set of movies used for all reconstructions.

**Tab. S2** List of all oligo nucleotides used in this study.

| Name | Sequence |
|---|---|
| pRS406_for_Orm1 | ATGACCGAATTAGATTATCAAGGAACTG |
| pRS406_rev_Orm1 | GGCCCTACGCGCTCTAGA |
| GAL1_for_OH_Orm1promoter | CTAGAGCGCGTAGGGCCAGTACGGATTAGAAGCCGC |
| GAL1_rev_OH_ORM1 | TAATCTAATTCGGTCATGTTTTTTCTCCTTGACGTTAAAG |
| pRS404_GAL1_for | TATATCTAGAACTAGTGGATCCCC |
| pRS404_GAL1_rev | GTTTTTTCTCCTTGACGTTAAAG |
| TSC3_OH_GAL1pr_for | CGTCAAGGAGAAAAAACATGACACAACATAAAAGCTCG |
| TSC3_OH_pRS404_rev | GGGGATCCACTAGTTCTAGATATATCTGTGACTCGGATATGGAG |
| pRS403_Sac1_for | ATGACAGGTCCAATAGTGTAC |
| pRS403_Sac1pr_rev | CGATCAGGACGTCAGGG |
| GAL1_OH_Sac1pr_for | CCCTGACGTCCTGATCGAGTACGGATTAGAAGCCG |
| GAL1_OH_Sac1_rev | CACTATTGGACCTGTCATGTTTTTTCTCCTTGACGTTAAAG |
| pRS405_Lcb2_for | ATGAGTACTCCTGCAAACTATACC |
| pRS405_Lcb2_rev | CACCCAATCACCGCGCTT |
| GAL1pr_OH_Lcb2pr_for | AAGCGCGGTGATTGGGTGAGTACGGATTAGAAGCCG |
| GAL1pr_OH_Lcb2_rev | TTTGCAGGAGTACTCATGTTTTTTCTCCTTGACGTTAAAG |
| pRS406_FLAG_LCB1_for | ATGGCACACATCCCAGAG |
| pRS406_FLAG_LCB1_rev | GACAGAGCAGTATGTGAGG |
| GAL1pr_OH_Lcb1pr_for | CCTCACATACTGCTCTGTCAGTACGGATTAGAAGCCG |
| GAL1pr_OH_Lcb1pr_rev | TCTGGGATGTGTGCCATGTTTTTTCTCCTTGACGTTAAAG |
| GAL1_for_OH_Orm2promoter | TAGAGGGTCCCCGGCATAGTACGGATTAGAAGCCGC |
| GAL1_rev_OH_Orm2 | TTAGTGCGGTCAATCATGTTTTTTCTCCTTGACGTTAAAG |
| pRS406_for_Orm2 | ATGATTGACCGCACTAAAAACGAATC |
| pRS406_rev_Orm2 | ATGCCGGGGACCCTCTAG |
| Orm2_AAA_exchange_DDD_for | cgacGTAATATCACATGTGGAACAG |
| Orm2_AAA_exchange_DDD_rev | tcgtcCCGTCTTCTCCTATGGTC |
| LCB1_S1 | GTT ATT TAT CCT TTT TTC TTC CTT CCC ACC CAA AAA AAA AAA GCA ATG CGT ACG CTG CAG GTC GAC |
| LCB1_S2 | ATA TAT ATG TGC GTG TGC ATA TAC TGG CTT TCT ATT TTT AAT CGA TGA ATT CGA GCT CG |
| ORM1_S2 | AAA ATA TAA ATA TAG CAA AAA CAT CTA GAT ACA AGA TTG AAA TAA ACT ATG TTC AAT CGA TGA ATT CGA GCT CG |
| ORM1_S1 | AAG CAG AGT TAT TCT TAT TTT GTA TTT CAT TGC ATT TTT ATC CAT TTA GTT AAT GCG TAC GCT GCA GGT CGA C |
| ORM2_S1 | GAA TTA ACG CAA GAC TAT ACC ATT ATA AAA ACG CAT AAG AAA CAG TTT CAT CAT GCG TAC GCT GCA GGT CGA C |

| | |
|---|---|
| ORM2_S2 | TATATATATATACATATATGCGTATAGGCAGAGCCAACTAg aattcgagctcgtttaaac |
| Q5_Lcb1-TMD1D_for | AAG CCA CAA CAG AAA AAG AGT CTT C |
| Q5_Lcb1-TMD1D_rev | TGG ATC GTC ATG ATG CGA TTT C |
| LCB1_SacI_F | cgcGAGCTCCCTCACATACTGCTCTGTCAATATGGC |
| LCB1_XhoI_R | gcgCTCGAGGCCTTGGTGTGCTGTTTTAACCTTG |
| LCB1-FLAG-F | GGCACACATCCCAGAGGTTTTACCCGACTACAAAGACCAT GACGGTGATT |
| LCB1-FLAG-R | AATCACCGTCATGGTCTTTGTAGTCGGGTAAAACCTCTGG GATGTGTGCC |
| FLAG-LCB1-F | CGATTACAAGGATGACGATGACAAGAAATCAATACCGATT CCGGCATTTATTGTTACC |
| FLAG-LCB1-R | GGTAACAATAAATGCCGGAATCGGTATTGATTTCTTGTCAT CGTCATCCTTGTAATCG |
| LCB1-TMD1-F | CAAGAAATCGCATAAGCCCAACCTATCGCCCCA |
| LCB1-TMD1-R | CGATAGGTTGGGCTTATGCGATTTCTTGATGTACGAAACG |
| ALFA-LCB1-F | TCCAGGTTGGAAGAGGAATTACGTCGTCGTTTGACCGAAc ccAAATCAATACCGATTCCGGCATTTATTGTTACC |
| ALFA-LCB1-R | TTCGGTCAAACGACGACGTAATTCCTCTTCCAACCTGGAG GGTAAAACCTCTGGGATGTGTGCC |
| ORM2_XbaI_F | cgcTCTAGAGGGTCCCCGGCATTGAGG |
| ORM2_EcoRI_R | gcgGAATTCCCCTAGAGGCAAGATTGTAGCTGAAGCTGG |
| ORM1_XhoI_F | cgcCTCGAGGCGCGTAGGGCCGCCAGCG |
| ORM1-SacI_R | cgcGAGCTCGAAGCAGTACGTGAAATAGTGC |
| ORM2_AAA_F | CCAGTGACCGACCATAGGAGAAGACGGGCAGCCGCCGTA ATATCACATGTGGAACAGGAAACC |
| ORM2_AAA_R | GGTTTCCTGTTCCACATGTGATATTACGGCGGCTGCCCGT CTTCTCCTATGGTCGGTCACTGG |
| ORM2_DDD_F | CCAGTGACCGACCATAGGAGAAGACGGGACGACGACGTA ATATCACATGTGGAACAGGAAACC |
| ORM2_DDD_R | GGTTTCCTGTTCCACATGTGATATTACGTCGTCGTCCCGT CTTCTCCTATGGTCGGTCACTGG |
| Orm1_AAA fw | CCTGTGAAAGATCATAGAAGAAGGCGTGCTGCCGCCATA ATTTCACATGTGGAACCGG |
| Orm1_AAA rev | CCGGTTCCACATGTGAAATTATGGCGGCAGCACGCCTTCT TCTATGATCTTTCACAGG |
| Orm1_DDD fw | CCTGTGAAAGATCATAGAAGAAGGCGTGATGA CGACATAATTTCACATGTGGAACCGG |
| Orm1_DDD rev | CCGGTTCCACATGTGAAATTATGTCGTCATCAC GCCTTCTTCTATGATCTTTCACAGG |
| LCB1-KO-F | CGTGTAGGGTTATTTATCCTTTTTTCTTCCTTCCCACCCAA AAAAAAAAAGCACGGATCCCCGGGTTAATTAA |
| LCB1-KO-F | GCGCATTCTCTGGGCGCCGTGCACCACGTCAATTTGACA GAGATGATGTTACGGAATTCGAGCTCGTTTAAAC |

**Tab. S3** List of used transitions for targeted lipidomics.

| Description | Q1 mass | Q2 mass | CE (V) | CXP (V) | DP (V) | EP (V) |
|---|---|---|---|---|---|---|
| Ceramide d17:1/24:0 | 636.629 | 249.8 | 47 | 13 | 120 | 10 |
| Phytoceramide 42:0;3 | 668.655 | 282.4 | 50 | 5 | 100 | 5 |
| Phytoceramide 42:0;4 | 684.650 | 282.4 | 50 | 5 | 100 | 5 |

| | | | | | | |
|---|---|---|---|---|---|---|
| Phytoceramide 42:0;5 | 700.645 | 282.4 | 50 | 5 | 100 | 5 |
| Phytoceramide 44:0;3 | 696.686 | 282.4 | 50 | 5 | 100 | 5 |
| Phytoceramide 44:0;4 | 712.681 | 282.4 | 50 | 5 | 100 | 5 |
| Phytoceramide 44:0;5 | 728.676 | 282.4 | 50 | 5 | 100 | 5 |
| Sphingosine d17:1 | 286.274 | 69 | 55 | 10 | 51 | 10 |
| 3-ketosphinganine 18:0 | 300.290 | 60 | 25 | 6 | 156 | 10 |
| Phytosphingosine 18:0 | 318.300 | 60 | 45 | 10 | 166 | 10 |
| Phytosphingosine 20:0 | 346.332 | 60 | 45 | 10 | 166 | 10 |
| Dihydrosphingosine 18:0 | 302.305 | 60 | 23 | 8 | 66 | 10 |
| Dihydrosphingosine 20:0 | 330.337 | 60 | 23 | 8 | 66 | 10 |

**Tab. S4** List of all exact *P*-values of LCB and ceramide analyses.

| **Measure** | | ***P*-value** |
|---|---|---|
| Total LCBS | ALFA-LCB1 – ALFA-lcb1TM1<br>ALFA-LCB1 – orm1Δ orm2Δ<br>ALFA-lcb1ΔTM1 – orm1Δ orm2Δ | 0.000189776780796<br>0.000000000545456<br>0.000000000549228 |
| Ceramide 44:0;4 | ALFA-LCB1 – ALFA-lcb1ΔTM1<br>ALFA-LCB1 – orm1Δ orm2Δ<br>ALFA-lcb1ΔTM1 – orm1Δ orm2Δ | 0.000593954054268<br>0.000000012538346<br>0.000000271128897 |
| Total LCBs+HS | ALFA-LCB1 – ALFA-lcb1ΔTM1<br>ALFA-LCB1 – orm1Δ orm2Δ<br>ALFA-lcb1ΔTM1 – orm1Δ orm2Δ | 0.000000000750627<br>0.000000000747786<br>0.997609131222421 |
| Ceramide 44:0;4+HS | ALFA-LCB1 – ALFA-lcb1ΔTM1<br>ALFA-LCB1 – orm1Δ orm2Δ<br>ALFA-lcb1ΔTM1 – orm1Δ orm2Δ | 0.723754341103371<br>0.009209620755693<br>0.030314697815936 |
| Total LCBs | orm2Δ ORM1 WT – orm2Δ ORM1 3A<br>orm1Δ ORM2 WT – orm1Δ ORM2 3A<br>orm2Δ ORM1 WT – orm1Δ ORM2 WT<br>orm2Δ ORM1 3A – orm1Δ ORM2 3A | 0.000347764131699<br>0.000000128982663<br>0.000311841874981<br>0.000000121051746 |
| Ceramide 44:0;4 | ORM2 3A – ORM1 3A<br>ORM2 3D – ORM1 3A<br>ORM2 3D – ORM2 3A | 0.083841240800197<br>0.179388138289236<br>0.826696422224932 |
